# Composite Likelihood Adjustment in Bayesian Inference for Estimating Species Tree Parameters Under the Multispecies Coalescent

**DOI:** 10.1101/2025.10.20.683470

**Authors:** Zixuan Chen, Laura S. Kubatko

## Abstract

Composite likelihood methods have long been used in biology to facilitate efficient computation for the high-dimensional, dependent structures commonly present in genetic data. In species-level phylogenetic inference under the multispecies coalescent model, the full likelihood is computationally intractable, while the composite likelihood can be efficiently computed. However, using the composite likelihood instead of the full likelihood in Bayesian inference involves repeated use of data, resulting in overly concentrated posteriors. We develop a method that incorporates an adjustment to the composite likelihood in a Bayesian framework to adjust the approximated posterior distributions so that credible intervals achieve the correct coverage. Both simulated and empirical data are used to show that our method compares favorably with existing methods with respect to both computation time and accuracy.

## 1 Introduction

Estimating the evolutionary relationship among a collection of present-day species is a problem of fundamental importance in the biological sciences. Such an evolutionary history is typically represented by a phylogenetic tree, a bifurcating, directed acyclic graph in which external vertices have degree one and internal vertices have degree three, except for the root vertex which has degree two. Time flows from the root vertex to the external vertices, which represent present-day species or populations generally referred to as taxa (singular: taxon). Molecular data are commonly used to infer the phylogenetic tree relating the taxa under consideration. As both the time and the cost required to obtain molecular data have decreased substantially in recent years, such data have become increasingly available for phylogenetic inference, with genome-scale data now in routine use. Such phylogenomic data present inference challenges because of the need to model the evolutionary process at the level of both genes and of species, each of which can be represented by a phylogenetic tree.

A phylogenetic *species tree* (*S*, ***τ, θ***) describes the evolutionary relationship among species, where *S* refers to the tree topology (branching pattern), *τ* is a vector of speciation times, and ***θ*** is a vector of effective population size parameters (the effective population size is defined to be 4*N μ*, where *N* is the census population size and *μ* is the mutation rate). In the example four-taxon species tree in Figure 1, the species tree topology *S* is drawn in black, the speciation times (*τ*_1_, *τ*_2_, *τ*_3_) of the species tree measure time backward from the present-day to the corresponding speciation event, and the population size parameters (*θ*_1_,…, *θ*_7_) correspond to the population sizes of the three ancestral populations (shaded in colors) and the four contemporary populations (unshaded). A *gene tree* (*G, t*) describes the evolutionary history among the gene copies sampled from individuals at a particular locus, where *G* refers to the tree topology, *t* is a vector of divergence times, and the term locus (plural: loci) refers to the physical position of a gene on a chromosome. Gene trees can be topologically incongruent with the species tree as the results of a variety of evolutionary processes (see, e.g., Maddison, 1997). Of these, the process of incomplete lineage sorting is widely accepted to generate variation in gene trees in empirical settings and is commonly modeled using the multispecies coalescent model (MSC) (Kingman, 1982b,a). A hypothetical gene tree that might have arisen under the MSC is depicted in blue in Figure 1. The nodes in the gene tree represent divergence events and (*t*_1_, *t*_2_, *t*_3_) measure the time interval between speciation events and gene divergence events (e.g., *t*_1_ and *t*_2_), or between consecutive gene divergence events (e.g., *t*_3_).

**Figure 1.**
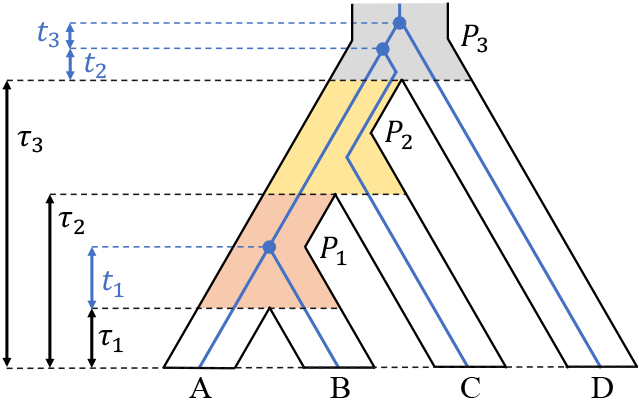
Example of a four-taxon species tree (in black) with a gene tree (in blue) nested within. The speciation times *τ*_1_, *τ*_2_, *τ*_3_ are measured backward from the present to the corresponding speciation event. The divergence times (*t*_1_, *t*_2_, *t*_3_) measure time between the most recent speciation event in the relevant time interval and the gene divergence event (i.e., *t*_1_ and *t*_2_) or between pairs of divergence events (i.e., *t*_3_). The ancestral populations (***P***_1_, ***P***_2_, ***P***_3_) are shaded in colors, whereas the four contemporary populations are unshaded and not labeled.

Computation of the likelihood of a species tree under the MSC model is complicated by the need to integrate over the large space of unobserved gene trees. Let ***D*** = (***D*** ^(1)^,…, ***D*** ^(*M*)^) be a collection of DNA matrices sampled at *M* loci. At locus *i*, let 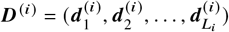) be the sample DNA data matrix, where the rows of ***D*** ^(*i*)^ are DNA sequences, the columns are aligned nucleotides, commonly known as sites, the vector 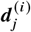 represents the nucleotides observed at site *j* in the alignment across taxa, and the length of ***D*** ^(*i*)^ is ***L***_*i*_. The likelihood of a species tree (*S, τ, θ*) under the MSC given data ***D*** is (Rannala, Yang, 2003)

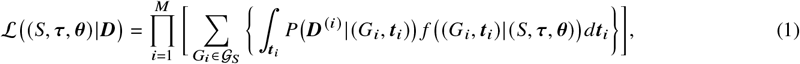

where 𝒢_*S*_ is the set of all possible gene tree topologies compatible with the species tree, *f* ((*G*_*i*_, ***t***_*i*_) | (*S*, ***τ, θ***)) is the density of the gene tree for locus *i* given species tree (*S, τ, θ*) under the MSC (see Rannala, Yang (2003) for details), and ***P*** (***D*** ^(*i*)^ | (*G*_*i*_, *t*_*i*_)) is the probability of the sequence data for gene *i* given the gene tree at locus *i*, which is computed via the Felsenstein likelihood (Felsenstein, 1981),

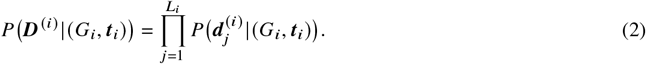

Unfortunately, several factors make the computation of the species tree likelihood in Equation 1 computationally intractable for large taxon samples. First, as the number of taxa grows, the number of tree topologies increases faster than exponentially, as shown in Table 1, and enumeration of possible topologies as required by the sum in Equation 1 is computationally daunting. Second, the integral over the gene tree divergence times has dimension equal to one less than the number of taxa, and thus also increases with the number of taxa. Finally, Equation 2 involves the product of very small site-wise conditional probabilities whose product is difficult to store with limitations on computer precision. Because this term appears within the integral in Equation 1, taking the logarithm does not help. For these reasons, computation of the species tree likelihood is currently limited to four-taxon trees with the additional assumption of site-wise independence within genes (see Methods for more details).

**Table 1.**
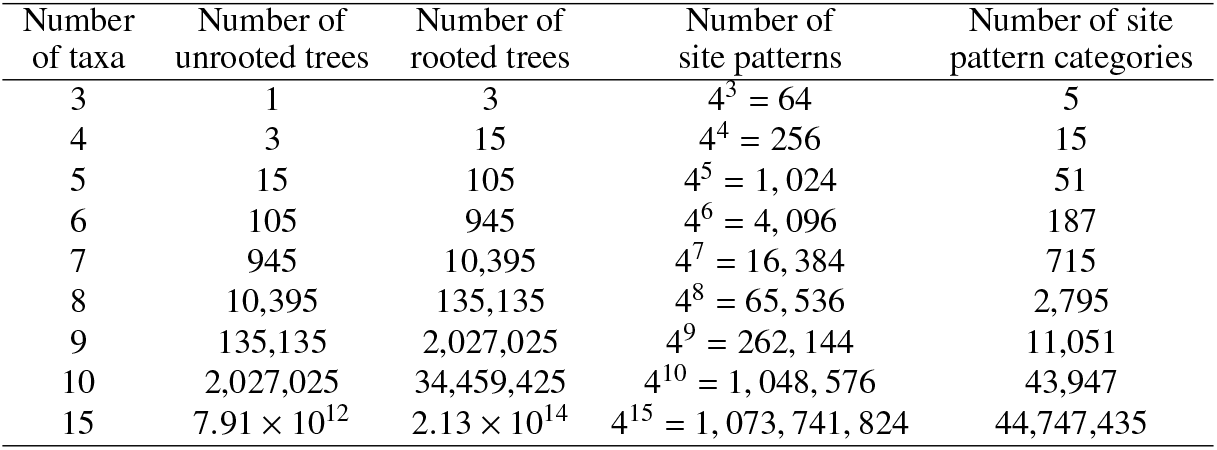
The number of possibilities for unrooted trees, rooted trees, site patterns, and site pattern categories for species trees with varying numbers of taxa.

| Number of taxa | Number of unrooted trees | Number of rooted trees | Number of site patterns | Number of site pattern categories |
| --- | --- | --- | --- | --- |
| 3 | 1 | 3 | $4^3 = 64$ | 5 |
| 4 | 3 | 15 | $4^4 = 256$ | 15 |
| 5 | 15 | 105 | $4^5 = 1,024$ | 51 |
| 6 | 105 | 945 | $4^6 = 4,096$ | 187 |
| 7 | 945 | 10,395 | $4^7 = 16,384$ | 715 |
| 8 | 10,395 | 135,135 | $4^8 = 65,536$ | 2,795 |
| 9 | 135,135 | 2,027,025 | $4^9 = 262,144$ | 11,051 |
| 10 | 2,027,025 | 34,459,425 | $4^{10} = 1,048,576$ | 43,947 |
| 15 | $7.91 \times 10^{12}$ | $2.13 \times 10^{14}$ | $4^{15} = 1,073,741,824$ | 44,747,435 |

Three classes of methods to circumvent the difficulty in computing the likelihood are currently used: 1) concatenation methods, 2) summary statistics methods (or two-step methods), and 3) full data methods. Concatenation methods combine DNA matrices from multiple loci into a supermatrix, thus assuming that all sites have the same gene tree, and estimate a tree using the gene tree likelihood in Equation 2 as their criterion. When the variation in gene trees from the MSC process is large, Kubatko, Degnan (2007) and Roch, Steel (2015) have shown that concatenation methods produce inconsistent estimates of the species tree with strong but incorrect bootstrap support.

Two-step methods first estimate gene trees through Equation 2 and then take the estimated gene trees as “data” to infer the species tree. They bypass the summation and integration in Equation 1 and use the conditional density *f* ((*G, t*) | (*S, τ, θ*)) as the optimization criterion. Two-step methods assume that the input gene trees are known without error; some have been proven to be statistically consistent under this assumption (see Roch, Warnow, 2015). However, in practice they are sensitive to errors in the input gene trees.

Full data methods are primarily Bayesian, utilizing Markov chain Monte Carlo (MCMC) to estimate a tree. They search the joint space of species trees and gene trees, drawing samples from the joint posterior with a specified prior for the species tree. The marginal posterior of the species tree *f* ((*S*, ***τ, θ***) | ***D***) is then estimated by considering the empirical marginal density of only the species trees from the MCMC draws. Full data methods have advantages in estimating species trees because their likelihood models the entire data-generating process for multilocus DNA data. However, full data methods are computationally intensive, often prohibitively so for datasets that are large in terms of either the number of species or the number of loci.

Composite likelihood provides an alternative approach to these three classes of methods. It has been used to obtain frequentist estimates of branch lengths for fixed species trees under the MSC model (e.g., Peng et al., 2022; Kubatko et al., 2025; Guerra, Nielsen, 2020). In particular, Peng et al. (2022) showed that composite likelihood estimates of branch lengths were consistent and asymptotically normal, and that such estimates could be obtained in a computationally efficient manner. Here, we consider extending this prior work to a Bayesian setting by using the composite likelihood within an MCMC algorithm.

When using composite likelihood in Bayesian analyses, the posterior sampled from the unadjusted composite likelihood is often overly concentrated because the composite likelihood treats overlapping events as independent and uses data repeatedly. Adjustments to the composite likelihood are thus necessary to obtain an appropriate approximation of the posterior distribution. Both magnitude adjustments (Pauli et al., 2011) and curvature adjustments (Ribatet et al., 2012) have been proposed to correct for this. The magnitude adjustment computes a positive constant that is multiplied with the composite log-likelihood and brings down the peak of the posterior to an appropriate magnitude. However, this adjustment can only be applied to composite likelihoods that are functions of a single parameter. When the composite likelihood is a function of a *p*-dimensional parameter, the curvature adjustment computes a *p* × *p* constant matrix ***C*** to transform the parameter space. The posteriors sampled from that parameter space have appropriate magnitude and curvature. The curvature adjustment was first proposed by Chandler, Bate (2007) in hypothesis testing and was later used by Ribatet et al. (2012) in Bayesian inference.

We propose a method that uses composite likelihood in MCMC for species-level phylogenetic inference by adapting the curvature adjustment of Ribatet et al. (2012) to our phylogenetic setting. We show that our method is more computationally efficient than existing full data methods while preserving accuracy. Section 2 introduces our method by defining the composite likelihood for species trees and providing details of the curvature adjustment. Section 3 shows the performance of our method in a simulation study and compares our method with another full data method in terms of efficiency and accuracy in an application to an empirical dataset. Section 4 concludes with a discussion of our findings, limitations of our method, and directions for future work.

## 2 Methods

### 2.1 Site pattern methods

In a DNA matrix with four rows (i.e., four DNA sequences), each column (i.e., site in the aligned data matrix) has 4^4^ = 256 possible patterns. Assuming the substitution model of Jukes, Cantor (1969), denoted JC69, the 256 site patterns can be grouped into 15 categories, where the site patterns in each category have the same probability of being observed under the model. These categories are

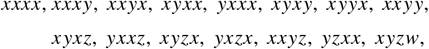

where *x, y, z*, and *w* denote the four distinct nucleotides (that is, *x* ≠ *y* ≠ *z* ≠ *w* ∈ {*A, C, G, T* }). For ease of notation, we denote the possible site patterns numerically by {1, 2,…, 15}. For example, site patterns such as {*AAAA, CCCC, GGGG, TTTT* } are sorted into category 1, {*AAAC, AAAG, AAAT*,…, *TTTG*} are sorted into category 2, and so on.

Our formulation of the species tree composite likelihood relies on consideration of the tree for four taxa at a time, which we call a quartet. Let (*Q, τ, θ*) denote a species quartet, with topology *Q*, vector of speciation times *τ*, and a single population size parameter *θ* for all populations, defined as in Figure 1. Focusing on only a single site (i.e., a locus of length 1 base pair (bp)), Chifman, Kubatko (2015) derived analytic expressions for the 15 site pattern probabilities of (*Q, τ, θ*) under the MSC for the JC69 substitution model. We denote these site pattern probabilities by *p*_*k*| (*Q*,***τ***,*θ*)_ for *k* = 1, 2,…, 15.

To compute the species quartet likelihood from the site pattern probabilities, we assume that all sites are unlinked unit-length loci under the MSC. Such data, called Coalescent Independent Sites (CIS), assume that every site is an independent and identically distributed sample from a multinomial distribution. The species quartet likelihood is thus a 15-category multinomial distribution where the category probabilities are the species quartet site pattern probabilities and the category counts are the corresponding site pattern counts observed in the aligned DNA sequence data, 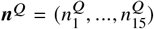 denote these observed quartet site pattern counts. Under these assumptions, the species quartet likelihood is defined as follows.

#### Definition 1 (Species quartet likelihood)

*Let* (*Q*, ***τ***, *θ*) *denote a four-taxon species tree where the population size parameter θ has the same value across populations. Let* {*p*_1| (*Q*,***τ***,*θ*)_,…, *p*_15| (*Q*,***τ***,*θ*)_ } *denote the species quartet site pattern probabilities and let* 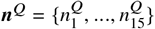 *denote the corresponding site pattern counts from the CIS data. The log likelihood of the species quartet is given by*

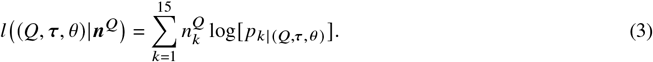

### 2.2 The composite likelihood of a species tree

Since the analytical form of the quartet site pattern probabilities has been derived by Chifman, Kubatko (2015), we can directly use the data to compute the species quartet likelihood in Equation 3 for quartets. The analytical form removes the need to integrate over gene trees, making the likelihood computation more efficient than that in Equation 1. However, for a species tree with more than four taxa, the full likelihood is intractable because its site pattern probabilities have not been derived analytically. Instead, a composite likelihood of the species tree can be formulated by enumerating all possible quartet subtrees and taking the product of the quartet subtree likelihoods.

#### Definition 2 (Species tree composite likelihood)

*Let* (*S*, ***τ***, *θ*) *denote an N-taxon species tree where the population size parameter θ has the same value across populations. Let* 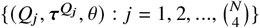 *denote the quartet subtrees of* (*S*, ***τ***, *θ*) *where* 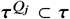 *is a sub-vector of speciation times for subtree Q*_*j*_. *Given CIS data, let* ***n***^*D*^ *denote the site pattern counts from the data, and let* 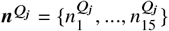 *denote the 15 category site pattern counts for subtree j*. *The composite log likelihood of the N-taxon species tree is the summation of the log likelihoods of the quartet subtrees*

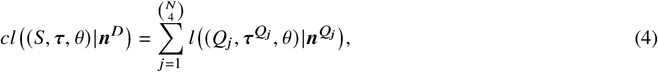

*where the log likelihood of a quartet subtree is given in Definition 1*.

In Figure 2, we give an example of a five-taxon species tree in the upper left, where the species are denoted by uppercase letters at the leaves. The full tree can be decomposed into 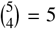 species quartet subtrees *Q*_1_,…, *Q*_5_ removing one species at a time, as indicated by the gray arrows. The speciation times of each quartet are a sub-vector of the full tree speciation times. For example, 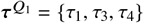 are the speciation times of the quartet *Q*_1_ and 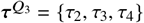 are the speciation times of the quartet *Q*_3_. Note that the quartets *Q*_1_ and *Q*_2_ have the same sub-vector of speciation times, implying that the species quartet site pattern probabilities of these two quartets have the same value under the model. Thus, their site pattern counts 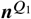 and 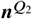 can be pooled when computing the composite likelihood. Likewise, when multiple quartet subtrees share the same sub-vector of speciation times, their site pattern counts can be pooled to save computation. In our method, the species tree topology *S* is fixed and we focus on estimating the vector of speciation times *τ* and the population size parameter *θ*. We use *cl* (***τ***, *θ* |***n***^*D*^) to denote the composite likelihood of (*τ, θ*) for a fixed species tree.

**Figure 2.**
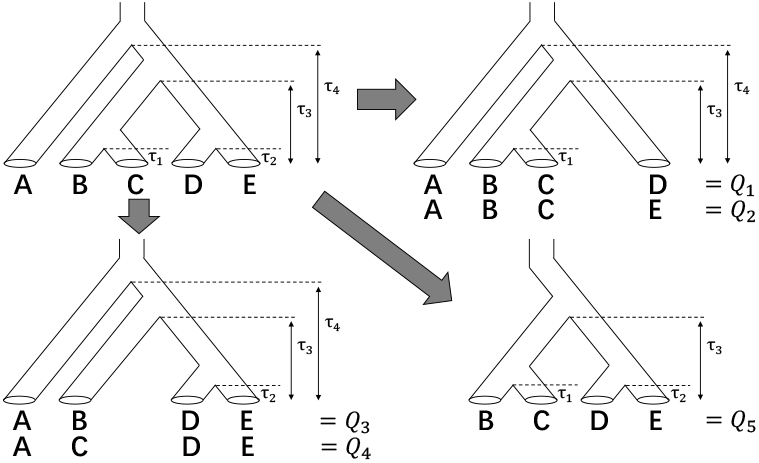
Decomposition of a species tree into quartet subtrees. A 5-taxon species tree (upper left) can be decomposed into 5 quartet subtrees: omission of species D or E leads to quartet subtrees *Q*_1_ or *Q*_2_ (upper right) with vectors of speciation times 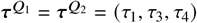; omission of species B or C leads to quartet subtrees *Q*_3_ or *Q*_4_ (lower left) with vectors of speciation times 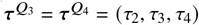; and omission of species A leads to quartet subtree *Q*_5_ (lower right) with vector of speciation times 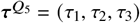.

### 2.3 Curvature adjustment

We choose the curvature adjustment of Ribatet et al. (2012), which we briefly review here, in our phylogenetic setting because our goal is to estimate the (*N* − 1)-dimensional vector of speciation times *τ*. For ease of notation, we use ***η*** = (***τ***, *θ*) to denote all the parameters of the species tree. The curvature-adjusted composite likelihood evaluates the composite likelihood at a translated, rescaled, and rotated parameter vector defined by

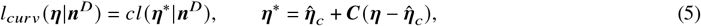

Where 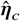 is the maximum composite likelihood estimator (MCLE) and ***C*** is a constant matrix computed from the sensitivity and variability matrices. The sensitivity matrix is given by 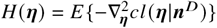 and the variability matrix is given by ***J*** (***η***) = *Var* {**∇**_***η***_ *cl* (***η***|***n***^*D*^)}, where **∇**_***η***_ *cl* (***η***|***n***^*D*^) denotes the gradient vector of the composite log-likelihood with respect to ***η***. Then, the curvature adjustment matrix is computed by ***C*** = ***M***^−1^ ***M***_*A*_ where

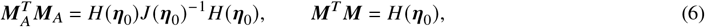

and ***η***_0_ denotes the true parameter value. Under some regularity conditions, Ribatet et al. (2012) showed that the curvature-adjusted composite posterior has the same asymptotic distribution as the MCLE 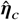. When we calculate the constant matrix ***C***, both ***H***(***η***_0_) and ***J*** (***η***_0_) must be estimated. In addition, the definition of matrix square root used to compute ***M*** and ***M***_*A*_ must also be defined, as it is not unique and can affect the performance of the curvature adjustment.

### 2.4 Computation of the curvature adjustment matrix in the phylogenetic setting

Estimation of the sensitivity and variability matrices involves deriving the first and second derivatives of the species quartet site pattern probabilities, and calculating the observed site pattern counts ***n***^*D*^ from the data. We have derived analytic expressions for the derivatives using Wolfram Mathematica and have included the formulas in our supplementary materials. The calculation of the observed site pattern counts ***n***^*D*^ depends on the size of the species tree. Note that the number of site pattern categories for an *N*-taxon tree is determined by 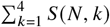, where *S*(*N, k*) denotes the Stirling number of the second kind (see, e.g., Richards, Kubatko, 2021), which grows very fast, as shown in Table 1. For large trees, enumerating all possible categories is cumbersome and unnecessary, since most categories will be unobserved in the data. Instead, we record the categories at their first occurrence in the data and accumulate counts at later occurrences. In practice, the number of site pattern categories observed in a given dataset is significantly less than the number of possible site pattern categories. For example, in a simulated dataset with 15 taxa and 1,000,000 sites, we recorded only 3,583 site pattern categories, compared to the 44,747,435 categories that are possible.

To calculate the site pattern counts ***n***^*D*^, we first need to differentiate the site pattern categories defined by the JC69 model, which we do by assigning each pattern a unique ID code. The details of this process are discussed in our supplementary materials. Then using the ID codes and ***n***^*D*^, we can get the 15 category site pattern counts of a quartet subtree *Q* by collapsing the site pattern counts ***n***^*D*^, that is ***n***^*Q*^ = ***E n***^*D*^, where ***E*** is a mapping matrix with 15 rows. Each row of ***E*** is a binary vector that has the same length as ***n***^*D*^ and sorts the selected counts of ***n***^*D*^ into the corresponding quartet site pattern counts by 15 categories.

To generate the binary vector in each row of ***E***, we need a mechanism that identifies the quartet site pattern (in *x, y, z*, and *w*) by its corresponding category numeric index. First, in a quartet site pattern of four nucleotides, we identify the nucleotides that do not repeat and count the number of unique nucleotides. For example, only the site pattern *xxxx* has one unique nucleotide, and only the site pattern *xyzw* has four unique nucleotides. We can identify them by category 1 and category 15, respectively, based on the count of unique nucleotides. However, the remaining thirteen site patterns will have either two or three unique nucleotides. Within each case, we need another rule to differentiate and identify each site pattern. If the count of unique nucleotides is two, we find the frequency of the most frequent nucleotide, which can be two or three. In the cases of three, the site patterns are *xxxy, xxyx, xyxx*, and *yxxx*, which correspond to categories 2, 3, 4, and 5, respectively. We observe that the string index of the least frequent nucleotide is identifiable for each site pattern, allowing us to establish a one-to-one mapping between them. Then, we can also sort the remaining site patterns to the corresponding category index; more details are provided in our supplementary materials.

For the performance of the curvature adjustment, we need an appropriate definition of the matrix square root for matrices ***M*** and ***M***_*A*_ to compute ***C*** = ***M***^−1^ ***M***_*A*_. However, the square root of a matrix is not unique because of rotational freedom. When using the curvature-adjusted composite likelihood for Bayesian inference, the rotation of the matrix square root determines the adjustment results of the sampled posteriors. In practice, if the matrix square root is not appropriately defined, some resulting posteriors can be overly flat and other posteriors can be overly concentrated. Referring to Miniato, Sartori (2021) and Kessy et al. (2018), we use a matrix square root definition called the zero-phase component analysis (ZCA). Similarly, Chandler, Bate (2007) have used this matrix square root definition for their adjustment but used it in hypothesis testing. They called it rotation-free adjustment since it preserves the structure of the parameter space when mapping from ***η*** to ***η***^***∗***^ in Equation 5. Let the eigen-decomposition of a square matrix **Σ** be **Σ** = ***U*Λ*U***^*T*^, with the eigenvector matrix ***U*** and the diagonal eigenvalue matrix **Λ**. The matrix square root of the ZCA for **Σ** is defined by

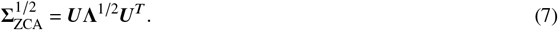

This definition does not alter the structure of the parameter space, as it first rotates via ***U***^*T*^, followed by scaling with **Λ**^1/2^, and then rotates back to the initial coordinate system using ***U***. As noted in Kessy et al. (2018), the square root of the inverse matrix can be computed by 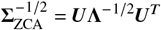.

### 2.5 MCMC algorithm

We use a multiple-block Metropolis algorithm (Algorithm 1) because the likelihood of the species tree is sensitive to small fluctuations in the population size parameter *θ*. We used two blocks to sequentially update the parameters of the species tree in a fixed order so that we can tune the step widths separately for the speciation times *τ* and the population size parameter *θ*.

In the MCMC, we use the unadjusted composite likelihood or the curvature-adjusted composite likelihood as an alternative likelihood to replace ℒ (*τ, θ*). The acceptance counters for *τ* and *θ* are used to calculate the acceptance ratios for tuning the step widths (*w*_*τ*_, *w*_*θ*_) so that the acceptance ratio of *τ* is around 23% and the acceptance ratio of *θ* is around 10%. Before sampling starts, we calculate the MCLE (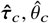) and use it as the initial value of the Markov chain, allowing us to bypass the burn-in period. We also use the MCLE as plug-in estimate in the computation of the curvature adjustment matrix ***C***. When proposed parameter values fail to satisfy the constraints imposed by the phylogeny, we propose new values until the constraints are satisfied.

#### Algorithm 1

Multiple-block Metropolis algorithm

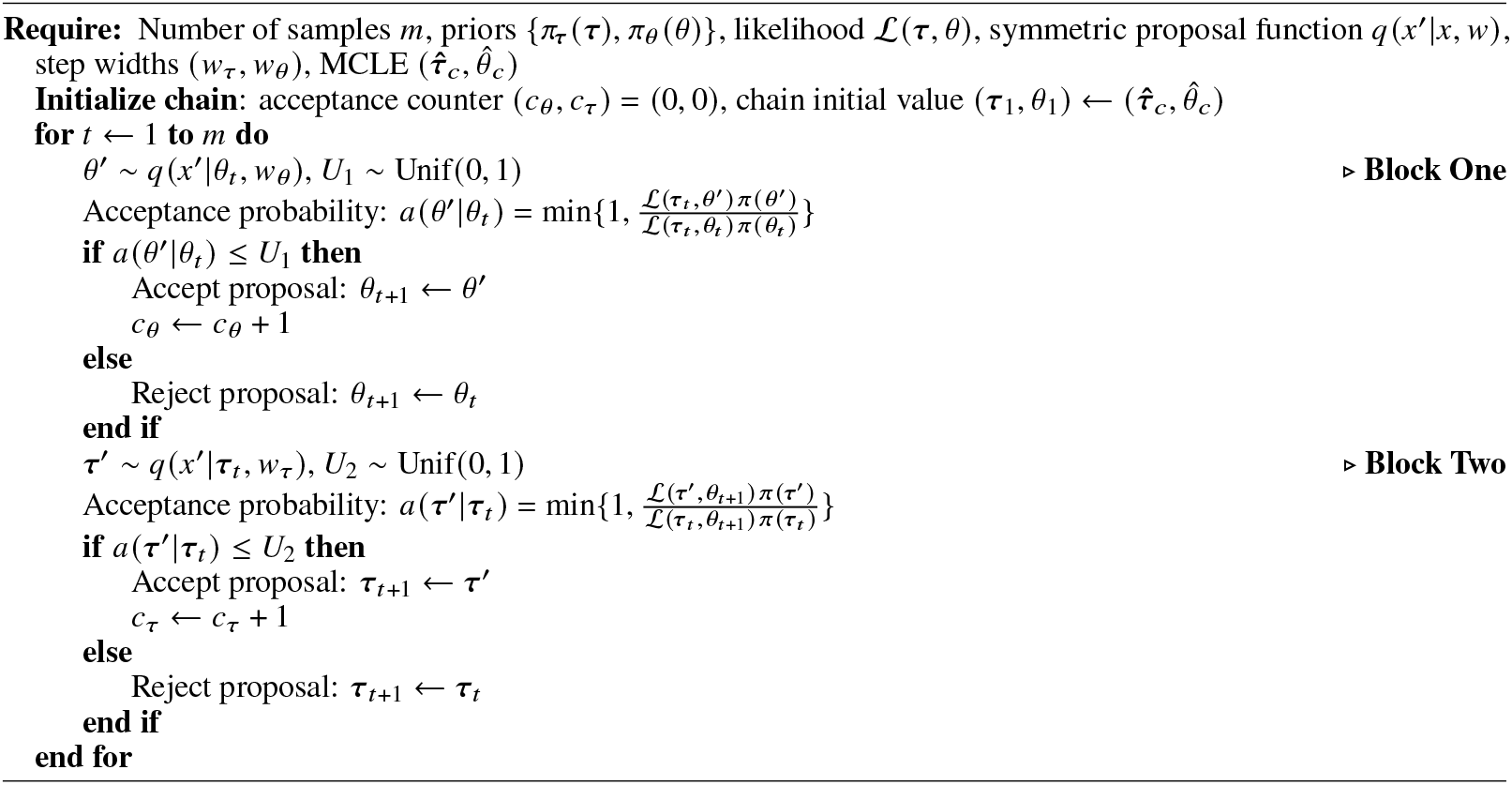

### 2.6 Prior distributions for the species tree parameters

Following Rannala et al. (2012), we use the inverse gamma distribution as the prior distribution for the population size parameter, i.e., we assume that *θ* ∼ *IG* (*α*_*θ*_, *β*_*θ*_). The hyperparameter *α*_*θ*_ > 0 determines the strength of our prior knowledge and *β*_*θ*_ > 0 determines the prior mean for a fixed *α*_*θ*_. For example, if we expect *θ* to be approximately 0.001, then the scale parameter can be set to *β*_*θ*_ = (*α*_*θ*_ − 1) × 0.001 (so that the prior mean is 0.001) and *α*_*θ*_ = 3 can be chosen to indicate weak prior information.

The prior distribution of the speciation times follows an inverse gamma compound Dirichlet distribution, as introduced by Rannala et al. (2012). In this prior specification, the root age of an *N*-taxon species tree follows the inverse gamma distribution *τ*_*N* −1_ ∼ *IG* (*α*_*τ*_, *β*_*τ*_), and the speciation times *τ*_1_,…, *τ*_*N* −2_ are assigned probability according to a flat Dirichlet distribution (details are given in the supplementary material). Hyperparameters for the inverse gamma distribution of the root age can be selected similarly to those for the population size parameter, as described above. Here, we give an example of the assignment of prior probability for the asymmetric quartet tree in Figure 1, where we have *τ*_3_ > *τ*_2_ > *τ*_1_. Given a fixed *τ*_3_, let 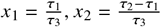 and 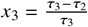 be the partition ratios that follow the flat Dirichlet distribution,

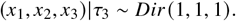

Applying the variable transformation (*x*_1_, *x*_2_) ↔ (*τ*_1_, *τ*_2_), we have

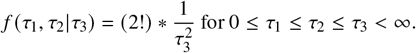

### 2.7 Simulation study

We evaluate the effectiveness of the curvature adjustment in MCMC by estimating the posterior distribution of the parameters for the 15-taxon species tree depicted in Figure 3 using both the unadjusted composite likelihood and the curvature-adjustment derived here. We assess performance using calibration plots that compare the nominal credible interval coverage with the empirical coverage in repeated simulation replicates. The steps in the simulation study are as follows.

**Figure 3.**
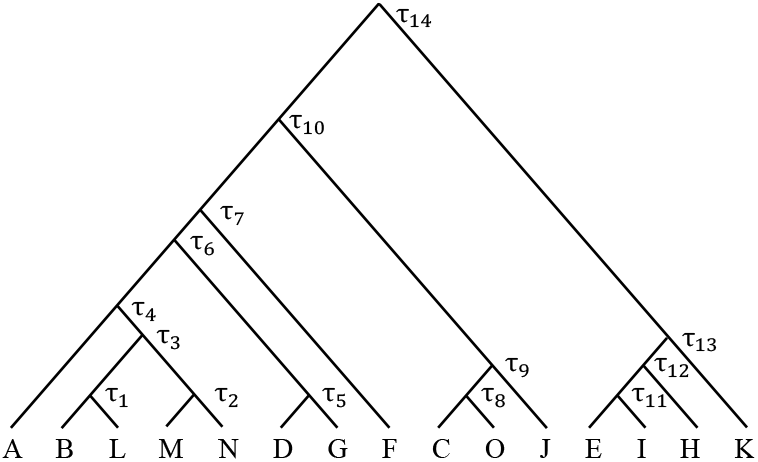
The 15-taxon species tree used in the simulation study. The capital letters at the tips represent present-day species. Speciation times (not drawn to scale) are labeled *τ*_1_,…, *τ*_14_.

1. Using PAUP^*^version 4a168 (Swofford et al., 2003), simulate 1,000,000 sites from the 15-taxon species tree in Figure 3 under the MSC with the JC69 substitution model.
2. Use the multiple-block Metropolis algorithm to draw posterior samples of (*τ, θ*) from the unadjusted composite likelihood *cl* (*τ, θ* |***n***^*D*^) and from the curvature-adjusted composite likelihoods *l*_*curv*_ (*τ, θ* |***n***^*D*^).
3. Draw posterior plots for (*τ, θ*) and construct equal-tailed credible intervals for each parameter at nominal levels ranging from 1% to 99%.
4. Repeat steps 1 to 3 for 200 repetitions and calculate the empirical coverage of the credible intervals at each nominal level.
5. Draw calibration plots for (*τ, θ*) comparing the nominal level and the empirical coverage for the approximate posterior distribution from the MCMC using both the unadjusted composite likelihood and the curvature-adjusted composite likelihood.

In step 1, CIS data were simulated along the 15-taxon species tree shown in Figure 3. The true parameter values are

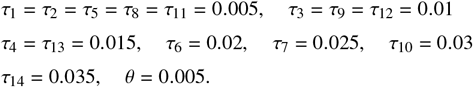

Speciation times are given in mutation units, and the population size parameter is used for all populations. Under the MSC, PAUP^*^simulated 1,000,000 gene trees within this species tree with a single DNA site for each gene tree (i.e., CIS data) using the JC69 substitution model.

In step 2, the means of the prior distributions for *τ*_14_ and *θ* were set to their MCLEs, and *α*_*θ*_ = 3 and *α*_*τ*_ = 3. MCMC samples were thinned by sampling every 100 iterations for the runs with both the unadjusted composite likelihood and the adjusted composite likelihood. For both approaches, sampling continued until 5,000 posterior draws were obtained for each parameter (i.e., a total of 500,000 iterations).

### 2.8 Assessing validity of the posterior distribution

To assess the performance of the curvature adjustment in composite likelihood, we need a definition of posterior validity (Monahan, Boos, 1992). Since the full likelihood is computationally inaccessible from the start, it is unrealistic to expect that the adjusted composite posterior can exactly match the posterior from the full likelihood. Instead, we anticipate that credible intervals constructed using the adjusted composite posteriors will have empirical coverage that is close to the nominal level. To assess this, we use the equal-tailed interval (ETI) in which the posterior probability below the left endpoint is equal to the posterior probability above the right endpoint. Using simulation, we summarize empirical coverage and draw calibration plots (Lazar, 2003; Miniato, Sartori, 2021) by plotting the empirical coverage on the *y* axis and the nominal credible level on the *x* axis. The ideal calibration curve is a 45-degree diagonal line from (0,0) to (1,1), indicating a perfect adjustment. However, in practice, the sample size is finite, and the results of the curvature adjustment may not be perfect. Nevertheless, we expect the curvature adjustment to bring the calibration curves close to the ideal 45-degree diagonal line.

### 2.9 Application to gibbon data

We explore the performance of our method in estimating speciation times on an empirical genome-scale dataset, which was earlier analyzed by Shi, Yang (2018), regarding five gibbon species: *Hylobates moloch* (Hm), *H. pileatus* (Hp), *Nomascus leucogenys* (N), *Hoolock leuconedys* (B), and *Symphalangus syndactylus* (S) (Carbone et al., 2014; Veeramah et al., 2015). The dataset consists of 11,323 coding loci with a length of 200 bp for each locus. The outgroup species (O) is human with a single lineage. The five gibbon species each have multiple lineages: N, B, and S each have four lineages, while Hm and Hp each have two. Fixing the species tree as shown in Figure 4, we use this dataset to estimate the branch lengths of the species tree (*b*_NBS_, *b*_BS_, and *b*_HpHm_) using both our method and *BPP* (Flouri et al., 2018), one of the state-of-the-art full data methods. These branch lengths are calculated by taking the difference between the two speciation times Δ*τ* associated with the corresponding branch.

**Figure 4.**
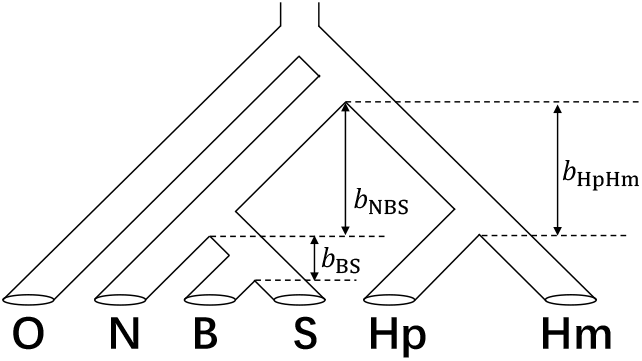
The species tree for the five gibbon species and the outgroup (O=human) with branch length parameters labeled by *b*_NBS_, *b*_BS_, and *b*_HpHm_. The subscripts in the branch length represent all the species descending from the lower node incident to the branch.

In contrast to our method, which samples the posterior distribution using a single value of the population size parameter *θ* across the entire tree, *BPP* simultaneously samples the posteriors of 10 population size parameters across all populations within the species tree. To compare the results of the three branch lengths in Figure 4 considering the difference between our model and that used in *BPP*, we convert the branch lengths from mutation units (number of per-site mutations given mutation rate *μ*) to coalescent units (number of generations scaled by the effective population size 2*N*). These two units can be inter-converted by *τ*_*coal*_ = (*θ*/2)*τ*_*mut*_ and *τ*_*mut*_ = (2/*θ*)*τ*_*coal*_.

Both our method and *BPP* use inverse gamma prior distributions for the root age and the population size parameters. We select *IG* (3, 0.002) as the prior distribution for the population size parameters and *IG* (3, 0.02) for the root age. Both methods used a thinning interval of 100 iterations. For *BPP*, we discarded the initial 5,000 iterations as burn-in. To compare the runtime of our method with *BPP*, we ran both methods on the same node of our computing cluster until 5,000 samples from the posterior were obtained. We also allowed *BPP* to continue running until the 2-week time limit of our cluster was reached.

Note that the sites in the gibbon data are not coalescent independent sites, which violates an assumption of our method. However, following Wascher, Kubatko (2021), we argue that our method is valid for both multilocus and CIS data as long as the number of loci is sufficiently large, as is the case for the gibbon data. Since the five gibbon species in this dataset include multiple lineages, we enumerate all possible quartet subtrees by selecting one lineage from each species, resulting in 752 quartets. We then pool the site pattern counts of the quartets that have the same sub-vector of speciation times, resulting in only 7 quartet likelihoods used to compute the composite likelihood, which significantly reduces the computational burden.

## 3 Results

### 3.1 Simulation study

Our method used about 7.5 hours to sample from the posteriors of the parameters for the 15-taxon species tree, with a range of 6.7 to 9 hours across the 200 simulation replicates. We could have further reduced the run time by running fewer MCMC iterations and requiring a smaller effective sample size (ESS). However, we were conservative in the sense that we required a larger ESS for all posterior samples to evaluate the posterior validity more precisely. Across repetitions of the simulation study, the ESS value ranged from roughly 300 to 800. Because we can numerically optimize the composite likelihood function to find the MCLE which is then used as the initial value, our method bypasses the burn-in period, reducing the required run time.

The posterior comparison plot in Figure 5 shows selected results from one of the 200 repetitions with comparisons between the posteriors sampled from the unadjusted composite likelihood (rawCL) and from the curvature-adjusted composite likelihood (adjCL). The unadjusted composite posteriors are more concentrated than the curvature-adjusted posteriors because the data are reused in the composite likelihood calculation for the quartet subtrees. The curvature-adjusted posteriors, as expected, have relatively flatter densities. When the unadjusted composite posteriors misplace their mode, as in the posterior comparison plot of *τ*_2_, the curvature-adjusted posteriors also behave similarly. However, the flatter densities of the curvature-adjusted posterior allow their credible intervals to have adequate coverage.

**Figure 5.**
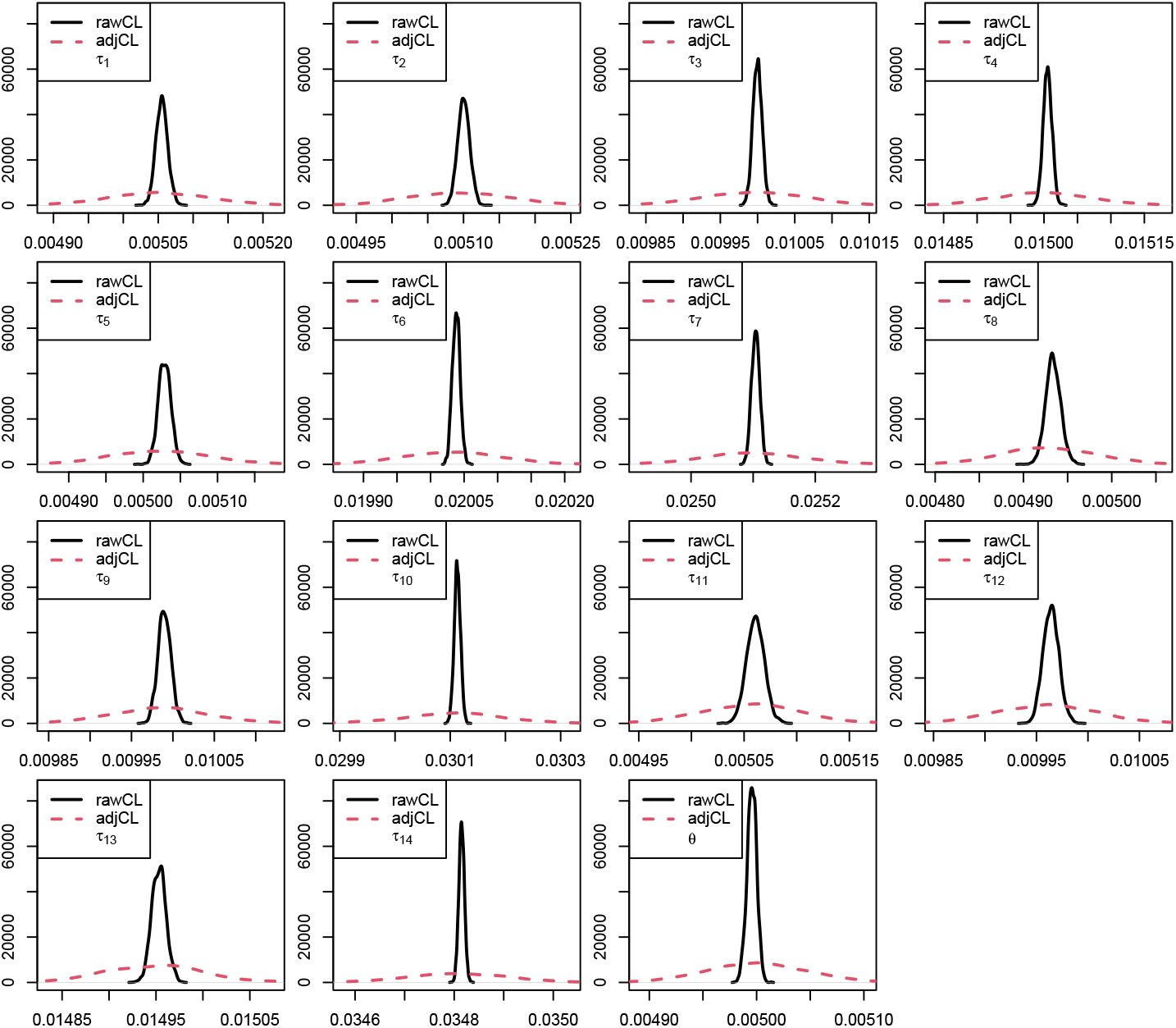
Marginal posteriors of (*τ*_1_,…, *τ*_14_, *θ*) from a single repetition of the simulation study with comparisons between the posteriors sampled from the unadjusted composite likelihood (rawCL) and from the curvature-adjusted composite likelihood (adjCL).

The calibration plots in Figure 6 were drawn by summarizing the empirical coverage from the 200 simulation replicates and plotting it against the nominal coverage. Because the unadjusted composite posteriors are more concentrated, their calibration curves are all well below the 45-degree diagonal line, implying that the over-concentration of the unadjusted composite posteriors leads to incorrect inference. In contrast, the calibration curves of the curvature-adjusted composite posteriors roughly match the 45-degree line. This demonstrates that the use of the curvature adjustment in Bayesian inference performs well in the phylogenetic setting.

**Figure 6.**
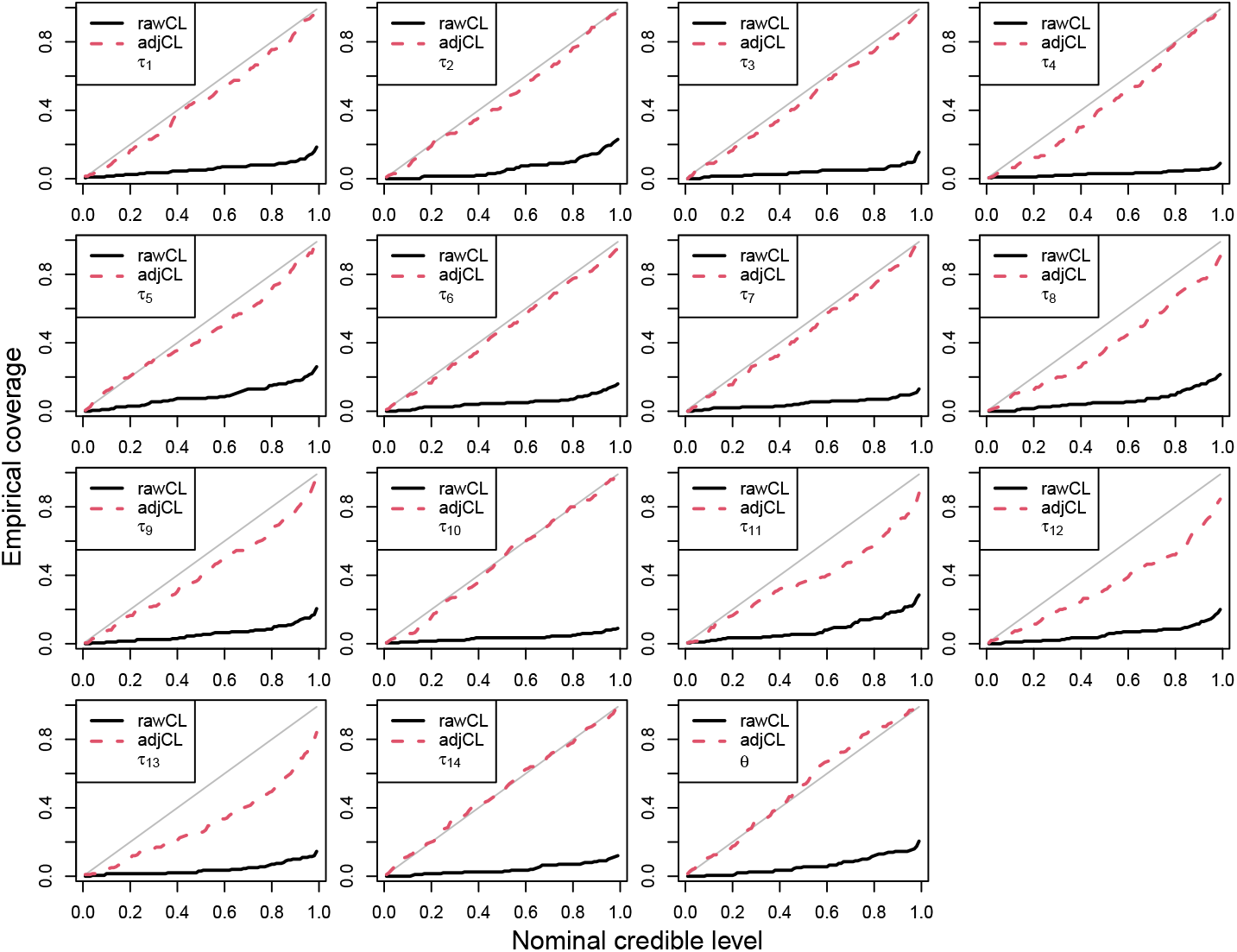
Calibration plots of (*τ*_1_,…, *τ*_14_, *θ*) computed from 200 repetitions of the simulation study with comparisons between the posteriors sampled from the unadjusted composite likelihood (rawCL) and from the curvature-adjusted composite likelihood (adjCL). The closer to the gray diagonal line the better.

### 3.2 Application to the gibbon data

Our method, which took approximately 33.3 minutes to complete, stored 5,000 posterior samples of six parameters, including five speciation times and a single population size parameter. In contrast, *BPP* used nearly two weeks to draw 18,759 posterior samples of 15 parameters, including 5 speciation times and 10 population size parameters. To compare the two methods, we converted from the speciation times measured in mutation units to the branch lengths measured in coalescent units.

Although 5,000 iterations were discarded as burn-in by *BPP* before sampling, we discarded an additional 2,000 samples (equivalent to 2, 000 × 100 = 200, 000 iterations) to attempt to achieve a thorough mixing of the chain. Trace plots for the *BPP* runs in Figure 7a and 7c indicate that the chains of both *b*_*NBS*_ and *b*_*BS*_ have poor mixing, showing evidence of a lack of convergence. Furthermore, as shown in Table 2, the ESS values of *b*_*NBS*_ and *b*_*BS*_ for the *BPP* runs are quite low, raising concerns about the reliability of the results from *BPP*. Trace plots for our method in Figure 7b, 7d, and 7f show good mixing with no evidence of lack of convergence. The ESS values of the three branch lengths for our method are all adequate as shown in Table 2.

**Table 2.** Means, 95% ETI in coalescent units, and ESS values for the three branch lengths *b*_NBS_, *b*_BS_, and *b*_HpHm_. The information in parentheses following the method name indicates the runtime used by each method. In our method, branch lengths are converted from mutation units to coalescent units using Δ*τ θ*, while for *BPP*, the conversion follows the formula in Shi, Yang (2018) (their Table 5).

| Branch | Our method<br>(33.3 minutes) |  |  | *BPP* - 5,000 samples<br>(125.2 hours) |  |  | *BPP* - 16,759 samples<br>(2 weeks) |  |  |
| --- | --- | --- | --- | --- | --- | --- | --- | --- | --- |
|  | Mean | 95% ETI | ESS | Mean | 95% ETI | ESS | Mean | 95% ETI | ESS |
| $b_{NBS}$ | 0.097 | (0.078, 0.116) | 4,385 | 0.067 | (0.047, 0.092) | 21 | 0.073 | (0.052, 0.094) | 71 |
| $b_{BS}$ | 0.051 | (0.038, 0.064) | 4,999 | 0.057 | (0.046, 0.069) | 33 | 0.053 | (0.039, 0.067) | 33 |
| $b_{HpHm}$ | 1.900 | (1.845, 1.957) | 2,497 | 2.056 | (1.974, 2.142) | 3,664 | 2.055 | (1.971, 2.141) | 10,700 |

**Figure 7.**
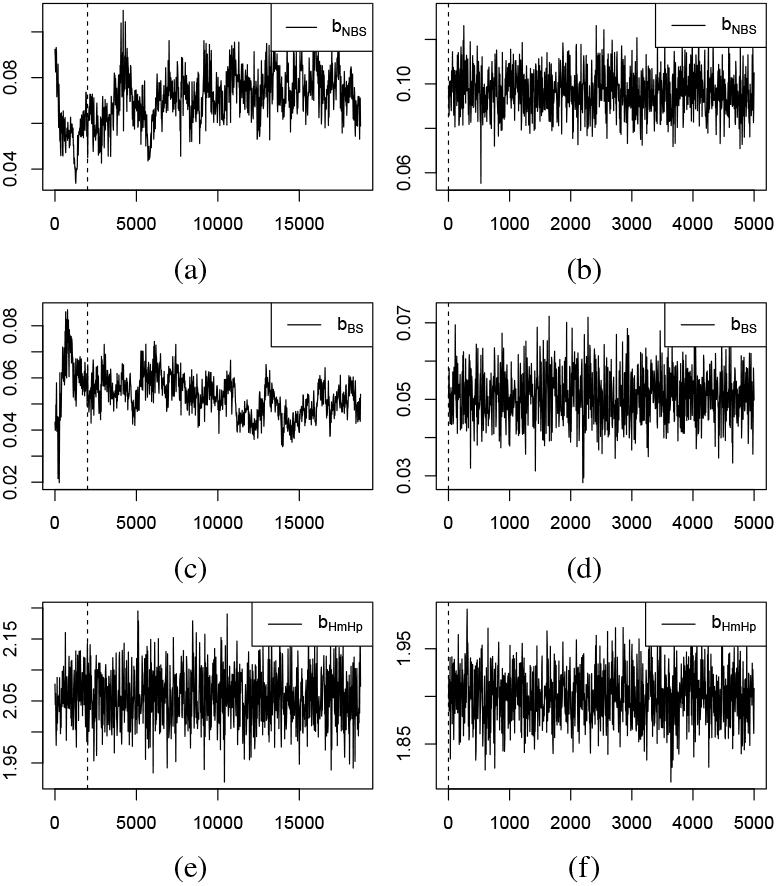
Trace plots of branch lengths *b*_NBS_, *b*_BS_, and *b*_HpHm_ for the method proposed here and for *BPP*. The first column (plots a, c, e) corresponds to the *BPP* runs whereas the second column (plots b, d, f) corresponds to our runs. Each row corresponds to branch lengths *b*_NBS_, *b*_BS_, and *b*_HpHm_ respectively. The dashed line at 2,000 in the *BPP* runs represents burn-in set to 2,000, whereas the dashed line at 0 in our runs represents no burn-in.

We compared the mean and 95% ETI of the posterior distributions for the three branch lengths in Table 2 between the two methods. Our method drew 5,000 posterior samples in 33.3 minutes, while *BPP*, under similar settings, took 125.2 hours to sample 5,000 posterior samples after discarding the first 2,000 as additional burn-in. The means of the three branch lengths for our method are similar in value to those of *BPP* with 5,000 samples. The credible intervals of *b*_*NBS*_ and *b*_*BS*_ overlap and have similar widths. However, the credible intervals of *b*_*HpHm*_ do not overlap. We also included the results for the *BPP* runs using all 16,759 posterior samples, which took about 2 weeks. The extended runtime included more samples, but it did not significantly increase the ESS for *b*_*NBS*_ and *b*_*BS*_, nor did it substantially impact the means or credible intervals.

## 4 Discussion

Inference of a phylogenetic tree from DNA sequence data is a challenging task that is crucially important in studying the evolutionary processes that lead to the formation of species. When data are collected at the level of genomes (i.e., data for many loci across the genome are available), the task is particularly challenging due to the need to model variation in the evolutionary histories of individual genes as well as variation in the DNA sequence data within the genes themselves. The MSC model is widely accepted as the state-of-the-art for capturing variability at both levels, but the likelihood for species trees with more than four species is computationally intractable under this model. To deal with this, we introduce a method that employs curvature-adjustment to the composite likelihood in MCMC to estimate the speciation times along a species tree under the MSC model. Our fully Bayesian approach has the potential to offer a significant efficiency boost while maintaining accuracy in species-level phylogenetic inference, as demonstrated using both simulation and an application to an empirical data set for gibbons.

To construct the composite likelihood based on the species quartet site pattern probabilities, we made three assumptions. First, we assumed a fixed value of the population size parameter *θ* across the species tree. While population size likely varies across species over evolutionary time, the configuration of available samples impacts whether the population size parameters and speciation times are both identifiable. In cases where species are closely related, such as the gibbon data set considered here, the assumption of a shared population size is likely to be a reasonable approximation.

Our method also assumes that the sequence data are generated via the CIS model. When the number of loci sampled is large (so that variation in gene trees is adequately represented in the sample), the quartet likelihood formed under the CIS model can itself be viewed as a composite likelihood, and the method can reasonably be expected to perform well. We do not recommend that our method be used when fewer than 50 loci are available for inference.

Finally, we assume that the sequence data evolve along the gene trees according to the JC69 substitution model. This was required so that the analytic expressions for the quartet site pattern probabilities derived by Chifman, Kubatko (2015) could be used in the quartet likelihood calculation. While derivation of corresponding analytic expressions under more complex substitution models is daunting, it may be possible to approximate the required substitution probabilities, and we leave this to future work.

Our current method estimates parameters along a fixed species tree topology. A logical extension is to develop an MCMC method that searches the space of species tree topologies as well. MCMC algorithms for searching tree space are well-know (e.g., Felsenstein, 2004), and our future work involves adapting current approaches to traverse the tree space and using composite likelihood in determination of the acceptance probability. However, unlike numerically continuous tree parameters, the tree topology is discrete and its space is vast, making the MCMC algorithm challenging to tune.

We also plan to extend our method to use different data types, such as single nucleotide polymorphism (SNP) data and data with site-to-site rate variation. These features have been implemented in the frequentist setting by Kubatko et al. (2025), where they have been found to perform well. In addition, the composite likelihood framework can be used to develop a hypothesis testing tool for species-level phylogenetic inference, as the curvature adjustment was first applied in hypothesis testing in Chandler, Bate (2007).

In summary, our method successfully integrates the composite likelihood curvature-adjustment for MCMC proposed by Ribatet et al. (2012) in the phylogenetic setting. The promising performance of the method was demonstrated with both simulated and empirical data. Incorporating composite likelihood with curvature adjustment has great potential for solving previously intractable problems in phylogenetic inference, and our method advances the state-of-the-art for parameter estimation on a fixed species tree under the multispecies coalescent.

