## Supplementary materials for "Composite Likelihood Adjustment in Bayesian Inference for Estimating Species Tree Parameters Under the Multispecies Coalescent"

### Supplementary materials to curvature-adjusted composite likelihood

Zixuan Chen

October 20, 2025

### A Gradient and Hessian of species quartet likelihood

#### A.1 Symmetric quartet

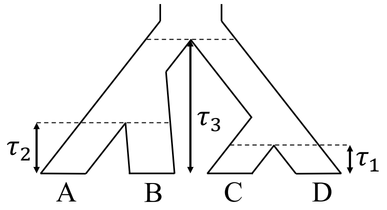


Figure 1: Symmetric quartet according to Chifman and Kubatko (2015)

There are 15 categories for quartet site pattern probabilities, $\{xxxx, xxxy, xxyx, xyxx, yxxx, xyxy, yxxy, xxyy, xyxz, xyzx, yxxz, yxzx, xxyz, yzxx, xyzw\}$, that can be further grouped into 9 bins:

| 1 | 2 | 3 | 4 | 5 | 6 | 7 | 8 | 9 |
|---|---|---|---|---|---|---|---|---|
| $xxxx$ | $xxxy$ | $xyxx$ | $xyxy$ | $xxyy$ | $xxyz$ | $yzxx$ | $xyxz$ | $xyzw$ |
| | $xxyx$ | $yxxx$ | $yxxy$ | | | | $yxxz$ | |
| | | | | | | | $yxzx$ | |
| | | | | | | | $xyzx$ | |

According to the supplement material of Chifman and Kubatko (2015), the vector of 15 category site pattern probabilities can be written as

$$\boldsymbol{p} \equiv \left(p^{\rho}_{1|(Q_s,\boldsymbol{\tau},\theta)}, p^{\rho}_{2|(Q_s,\boldsymbol{\tau},\theta)}, ..., p^{\rho}_{15|(Q_s,\boldsymbol{\tau},\theta)}\right)^T = \boldsymbol{W}_s \boldsymbol{C}_s^T \boldsymbol{\beta}_s = \boldsymbol{W}_s(\boldsymbol{c}_{s1}^T\boldsymbol{\beta}_s, \boldsymbol{c}_{s2}^T\boldsymbol{\beta}_s, ..., \boldsymbol{c}_{s9}^T\boldsymbol{\beta}_s)^T,$$

where 1) $Q_s$ is the symmetric quartet topology, 2) $\boldsymbol{W}_s$ is a transformation weight matrix

$$\boldsymbol{W}_s = \begin{pmatrix} 4 & 0 & 0 & 0 & 0 & 0 & 0 & 0 & 0 \\ 0 & 12 & 0 & 0 & 0 & 0 & 0 & 0 & 0 \\ 0 & 12 & 0 & 0 & 0 & 0 & 0 & 0 & 0 \\ 0 & 0 & 12 & 0 & 0 & 0 & 0 & 0 & 0 \\ 0 & 0 & 12 & 0 & 0 & 0 & 0 & 0 & 0 \\ 0 & 0 & 0 & 12 & 0 & 0 & 0 & 0 & 0 \\ 0 & 0 & 0 & 12 & 0 & 0 & 0 & 0 & 0 \\ 0 & 0 & 0 & 0 & 12 & 0 & 0 & 0 & 0 \\ 0 & 0 & 0 & 0 & 0 & 0 & 0 & 24 & 0 \\ 0 & 0 & 0 & 0 & 0 & 0 & 0 & 24 & 0 \\ 0 & 0 & 0 & 0 & 0 & 0 & 0 & 24 & 0 \\ 0 & 0 & 0 & 0 & 0 & 0 & 0 & 24 & 0 \\ 0 & 0 & 0 & 0 & 0 & 24 & 0 & 0 & 0 \\ 0 & 0 & 0 & 0 & 0 & 0 & 24 & 0 & 0 \\ 0 & 0 & 0 & 0 & 0 & 0 & 0 & 0 & 24 \end{pmatrix},$$

3) $\boldsymbol{C}_s = (\boldsymbol{c}_{s1}, \boldsymbol{c}_{s2}, ..., \boldsymbol{c}_{s9})$ is a $9 \times 9$ coefficient matrix, 4) $\boldsymbol{\beta}_s = \left(1, x_1^{2\rho\mu}, x_2^{2\rho\mu}, x_1^{2\rho\mu}x_2^{2\rho\mu}, x_3^{2\rho\mu}, x_1^{\rho\mu}x_3^{2\rho\mu}, x_2^{\rho\mu}x_3^{2\rho\mu}, x_1^{\rho\mu}x_2^{\rho\mu}x_3^{2\rho\mu}, x_1^{-\frac{2}{\theta}}x_2^{-\frac{2}{\theta}}x_3^{4(\rho\mu+\frac{1}{\theta})}\right)^T$ is a $9 \times 1$ parameter vector, 5) $x_i = e^{-\tau_i}$ for $i \in \{1,2,3\}$ and 6) $\rho$ is the various evolution rate.

In our algorithm, the user input population size parameter $\tilde{\theta}$ is converted to the default $\theta$ of Chifman and Kubatko (2015) by $\theta = 2\tilde{\theta}$. Since our MCMC algorithm samples posterior for $\tilde{\theta}$, we take derivative with respect to $\tilde{\theta}$, which by chain rule gives us $\frac{\partial\theta}{\partial\tilde{\theta}} = 2$. For convenience of notation, we use $\odot$ to denote the Hadamard product (i.e., element-wise product) and $\oslash$ to denote the Hadamard division (i.e., element-wise division) between vectors. Fixing the species quartet topology $Q_s$, the log-likelihood of the symmetric species quartet given the quartet site pattern counts is denoted by

$$l(\boldsymbol{\tau}, \theta | \boldsymbol{n}^Q) = \sum_{k=1}^{15} n_k^Q \log(p_k) = (\boldsymbol{n}^Q)^T \log(\boldsymbol{p}). \tag{1}$$

##### A.1.1 First derivatives

We derive the first-order partial derivatives of the log likelihood $\left(\frac{\partial}{\partial\tau_1}l(\boldsymbol{\tau},\theta), \frac{\partial}{\partial\tau_2}l(\boldsymbol{\tau},\theta), \frac{\partial}{\partial\tau_3}l(\boldsymbol{\tau},\theta), \frac{\partial}{\partial\tilde{\theta}}l(\boldsymbol{\tau},\theta)\right)^T$ where

$$\frac{\partial}{\partial\tau_i}l(\boldsymbol{\tau},\theta) = \sum_{k=1}^{15}\frac{n_k^Q}{p_k}\cdot\frac{\partial}{\partial\tau_i}p_k = \left(\boldsymbol{n}^Q \oslash \boldsymbol{p}\right)^T\left(\frac{\partial}{\partial\tau_i}\boldsymbol{p}\right) \text{ for } i = 1,2,3,$$
$$\frac{\partial}{\partial\theta}l(\boldsymbol{\tau},\theta) = \sum_{k=1}^{15}\frac{n_k^Q}{p_k}\cdot\frac{\partial}{\partial\theta}p_k = \left(\boldsymbol{n}^Q \oslash \boldsymbol{p}\right)^T\left(\frac{\partial}{\partial\theta}\boldsymbol{p}\right).$$

The first-order partial derivatives of the site pattern probabilities are derived as follows.

**First-order partial derivatives for $\tau_i$**

$$\frac{\partial}{\partial\tau_i}\boldsymbol{p} = \boldsymbol{W}_s\boldsymbol{C}_s^T\left(\frac{\partial}{\partial\tau_i}\boldsymbol{\beta}_s\right) = \boldsymbol{W}_s\boldsymbol{C}_s^T\left(\gamma_{\tau_i}\odot\boldsymbol{\beta}_s\right) \text{ for } i = 1,2,3,$$

where

- $\gamma_{\tau_1} = \left(0, -2\rho\mu, 0, -2\rho\mu, 0, -\rho\mu, 0, -\rho\mu, \frac{2}{\theta}\right)^T$,
- $\gamma_{\tau_2} = \left(0, 0, -2\rho\mu, -2\rho\mu, 0, 0, -\rho\mu, -\rho\mu, \frac{2}{\theta}\right)^T$, and
- $\gamma_{\tau_3} = \left(0, 0, 0, 0, -2\rho\mu, -2\rho\mu, -2\rho\mu, -2\rho\mu, -4(\rho\mu + \frac{1}{\theta})\right)^T$.

The partial derivatives in parentheses are represented as the Hadamard product $\gamma_{\tau_i}\odot\boldsymbol{\beta}_s$ because all the speciation times $\tau_i$ are part of exponential terms.

**First-order partial derivative for $\tilde{\theta}$**

$$\begin{aligned}\frac{\partial}{\partial\tilde{\theta}}\boldsymbol{p} &= \boldsymbol{W}_s\left[\boldsymbol{C}_s^T\left(\frac{\partial}{\partial\tilde{\theta}}\boldsymbol{\beta}_s\right) + \left(\frac{\partial}{\partial\tilde{\theta}}\boldsymbol{C}_s\right)^T\boldsymbol{\beta}_s\right] \\ &\overset{\partial\theta=2\partial\tilde{\theta}}{=} \boldsymbol{W}_s\left[\boldsymbol{C}_s^T\left(2\frac{\partial}{\partial\theta}\boldsymbol{\beta}_s\right) + \left(2\frac{\partial}{\partial\theta}\boldsymbol{C}_s\right)^T\boldsymbol{\beta}_s\right] \\ &= \boldsymbol{W}_s\left[2\boldsymbol{C}_s^T\left(\gamma_\theta\odot\boldsymbol{\beta}_s\right) + \left(2\frac{\partial}{\partial\theta}\boldsymbol{C}_s\right)^T\boldsymbol{\beta}_s\right]\end{aligned}$$

where $\gamma_\theta = \left(0,0,0,0,0,0,0,0,\frac{4\tau_3-2(\tau_1+\tau_2)}{\theta^2}\right)^T$, and $\left(\frac{\partial}{\partial\theta}\boldsymbol{C}_s\right) = \left(\frac{\partial}{\partial\theta}\boldsymbol{c}_{s1}, \frac{\partial}{\partial\theta}\boldsymbol{c}_{s2}, ..., \frac{\partial}{\partial\theta}\boldsymbol{c}_{s9}\right)$ is included in table 1. Some derivatives in table 1 are shown separately as they are too long for the table.

Table 1: First-order partial derivatives of $\theta$ to the symmetric quartet coefficient matrix $\boldsymbol{C}_s$

| $\frac{\partial}{\partial\theta}\boldsymbol{c}_{s1}$ | $\frac{\partial}{\partial\theta}\boldsymbol{c}_{s2}$ | $\frac{\partial}{\partial\theta}\boldsymbol{c}_{s3}$ |
|---|---|---|
| $0$ | $0$ | $0$ |
| $-\frac{(\kappa-1)\rho\mu}{\kappa^4(1+\rho\mu\theta)^2}$ | $\frac{\rho\mu}{\kappa^4(1+\rho\mu\theta)^2}$ | $-\frac{(\kappa-1)\rho\mu}{\kappa^4(1+\rho\mu\theta)^2}$ |
| $-\frac{(\kappa-1)\rho\mu}{\kappa^4(1+\rho\mu\theta)^2}$ | $-\frac{(\kappa-1)\rho\mu}{\kappa^4(1+\rho\mu\theta)^2}$ | $\frac{\rho\mu}{\kappa^4(1+\rho\mu\theta)^2}$ |
| $-\frac{2(\kappa-1)^2\rho\mu}{\kappa^4(1+\rho\mu\theta)^3}$ | $\frac{2(\kappa-1)\rho\mu}{\kappa^4(1+\rho\mu\theta)^3}$ | $\frac{2(\kappa-1)\rho\mu}{\kappa^4(1+\rho\mu\theta)^3}$ |
| $-\frac{4(\kappa-1)\rho\mu}{\kappa^4(1+\rho\mu\theta)^2}$ | $-\frac{2(\kappa-2)\rho\mu}{\kappa^4(1+\rho\mu\theta)^2}$ | $-\frac{2(\kappa-2)\rho\mu}{\kappa^4(1+\rho\mu\theta)^2}$ |
| $-\frac{4(\kappa-2)(\kappa-1)\rho\mu(2\rho\mu\theta+3)}{\kappa^4(1+\rho\mu\theta)^2(2+\rho\mu\theta)^2}$ | $\frac{4(\kappa-2)\rho\mu(2\rho\mu\theta+3)}{\kappa^4(1+\rho\mu\theta)^2(2+\rho\mu\theta)^2}$ | $-\frac{2(\kappa-2)^2\rho\mu(2\rho\mu\theta+3)}{\kappa^4(1+\rho\mu\theta)^2(2+\rho\mu\theta)^2}$ |
| $-\frac{4(\kappa-2)(\kappa-1)\rho\mu(2\rho\mu\theta+3)}{\kappa^4(1+\rho\mu\theta)^2(2+\rho\mu\theta)^2}$ | $-\frac{2(\kappa-2)^2\rho\mu(2\rho\mu\theta+3)}{\kappa^4(1+\rho\mu\theta)^2(2+\rho\mu\theta)^2}$ | $\frac{4(\kappa-2)\rho\mu(2\rho\mu\theta+3)}{\kappa^4(1+\rho\mu\theta)^2(2+\rho\mu\theta)^2}$ |
| $-\frac{4(\kappa-2)^2(\kappa-1)\rho\mu(3\rho\mu\theta+4)}{\kappa^4(1+\rho\mu\theta)^2(2+\rho\mu\theta)^3}$ | $\frac{4(\kappa-2)^2\rho\mu(3\rho\mu\theta+4)}{\kappa^4(1+\rho\mu\theta)^2(2+\rho\mu\theta)^3}$ | $\frac{4(\kappa-2)^2\rho\mu(3\rho\mu\theta+4)}{\kappa^4(1+\rho\mu\theta)^2(2+\rho\mu\theta)^3}$ |
| See $\frac{\partial}{\partial\theta}c_{8,1}$ | See $\frac{\partial}{\partial\theta}c_{8,2}$ | See $\frac{\partial}{\partial\theta}c_{8,3}$ |

| $\frac{\partial}{\partial\theta}\boldsymbol{c}_{s4}$ | $\frac{\partial}{\partial\theta}\boldsymbol{c}_{s5}$ | $\frac{\partial}{\partial\theta}\boldsymbol{c}_{s6}$ |
|---|---|---|
| $0$ | $0$ | $0$ |
| $\frac{\rho\mu}{\kappa^4(1+\rho\mu\theta)^2}$ | $-\frac{(\kappa-1)\rho\mu}{\kappa^4(1+\rho\mu\theta)^2}$ | $\frac{\rho\mu}{\kappa^4(1+\rho\mu\theta)^2}$ |
| $\frac{\rho\mu}{\kappa^4(1+\rho\mu\theta)^2}$ | $-\frac{(\kappa-1)\rho\mu}{\kappa^4(1+\rho\mu\theta)^2}$ | $-\frac{(\kappa-1)\rho\mu}{\kappa^4(1+\rho\mu\theta)^2}$ |
| $-\frac{2\rho\mu}{\kappa^4(1+\rho\mu\theta)^3}$ | $-\frac{2(\kappa-1)^2\rho\mu}{\kappa^4(1+\rho\mu\theta)^3}$ | $\frac{2(\kappa-1)\rho\mu}{\kappa^4(1+\rho\mu\theta)^3}$ |
| $-\frac{2(\kappa-2)\rho\mu}{\kappa^4(1+\rho\mu\theta)^2}$ | $\frac{4\rho\mu}{\kappa^4(1+\rho\mu\theta)^2}$ | $\frac{4\rho\mu}{\kappa^4(1+\rho\mu\theta)^2}$ |
| $\frac{4(\kappa-2)\rho\mu(2\rho\mu\theta+3)}{\kappa^4(1+\rho\mu\theta)^2(2+\rho\mu\theta)^2}$ | $\frac{4(\kappa-2)\rho\mu(2\rho\mu\theta+3)}{\kappa^4(1+\rho\mu\theta)^2(2+\rho\mu\theta)^2}$ | $-\frac{8\rho\mu(2\rho\mu\theta+3)}{\kappa^4(1+\rho\mu\theta)^2(2+\rho\mu\theta)^2}$ |
| $\frac{4(\kappa-2)\rho\mu(2\rho\mu\theta+3)}{\kappa^4(1+\rho\mu\theta)^2(2+\rho\mu\theta)^2}$ | $\frac{4(\kappa-2)\rho\mu(2\rho\mu\theta+3)}{\kappa^4(1+\rho\mu\theta)^2(2+\rho\mu\theta)^2}$ | $\frac{4(\kappa-2)\rho\mu(2\rho\mu\theta+3)}{\kappa^4(1+\rho\mu\theta)^2(2+\rho\mu\theta)^2}$ |
| $-\frac{8(\kappa-2)\rho\mu(3\rho\mu\theta+4)}{\kappa^4(1+\rho\mu\theta)^2(2+\rho\mu\theta)^3}$ | $\frac{4(\kappa-2)^2\rho\mu(3\rho\mu\theta+4)}{\kappa^4(1+\rho\mu\theta)^2(2+\rho\mu\theta)^3}$ | $-\frac{8(\kappa-2)\rho\mu(3\rho\mu\theta+4)}{\kappa^4(1+\rho\mu\theta)^2(2+\rho\mu\theta)^3}$ |
| See $\frac{\partial}{\partial\theta}c_{8,4}$ | See $\frac{\partial}{\partial\theta}c_{8,5}$ | See $\frac{\partial}{\partial\theta}c_{8,6}$ |

| $\frac{\partial}{\partial\theta}\boldsymbol{c}_{s7}$ | $\frac{\partial}{\partial\theta}\boldsymbol{c}_{s8}$ | $\frac{\partial}{\partial\theta}\boldsymbol{c}_{s9}$ |
|---|---|---|
| $0$ | $0$ | $0$ |
| $-\frac{(\kappa-1)\rho\mu}{\kappa^4(1+\rho\mu\theta)^2}$ | $\frac{\rho\mu}{\kappa^4(1+\rho\mu\theta)^2}$ | $\frac{\rho\mu}{\kappa^4(1+\rho\mu\theta)^2}$ |
| $\frac{\rho\mu}{\kappa^4(1+\rho\mu\theta)^2}$ | $\frac{\rho\mu}{\kappa^4(1+\rho\mu\theta)^2}$ | $\frac{\rho\mu}{\kappa^4(1+\rho\mu\theta)^2}$ |
| $\frac{2(\kappa-1)\rho\mu}{\kappa^4(1+\rho\mu\theta)^3}$ | $-\frac{2\rho\mu}{\kappa^4(1+\rho\mu\theta)^3}$ | $-\frac{2\rho\mu}{\kappa^4(1+\rho\mu\theta)^3}$ |
| $\frac{4\rho\mu}{\kappa^4(1+\rho\mu\theta)^2}$ | $-\frac{(\kappa-4)\rho\mu}{\kappa^4(1+\rho\mu\theta)^2}$ | $\frac{4\rho\mu}{\kappa^4(1+\rho\mu\theta)^2}$ |
| $\frac{4(\kappa-2)\rho\mu(2\rho\mu\theta+3)}{\kappa^4(1+\rho\mu\theta)^2(2+\rho\mu\theta)^2}$ | $\frac{2(\kappa-4)\rho\mu(2\rho\mu\theta+3)}{\kappa^4(1+\rho\mu\theta)^2(2+\rho\mu\theta)^2}$ | $-\frac{8\rho\mu(2\rho\mu\theta+3)}{\kappa^4(1+\rho\mu\theta)^2(2+\rho\mu\theta)^2}$ |
| $-\frac{8\rho\mu(2\rho\mu\theta+3)}{\kappa^4(1+\rho\mu\theta)^2(2+\rho\mu\theta)^2}$ | $\frac{2(\kappa-4)\rho\mu(2\rho\mu\theta+3)}{\kappa^4(1+\rho\mu\theta)^2(2+\rho\mu\theta)^2}$ | $-\frac{8\rho\mu(2\rho\mu\theta+3)}{\kappa^4(1+\rho\mu\theta)^2(2+\rho\mu\theta)^2}$ |
| $-\frac{8(\kappa-2)\rho\mu(3\rho\mu\theta+4)}{\kappa^4(1+\rho\mu\theta)^2(2+\rho\mu\theta)^3}$ | $-\frac{4(\kappa-4)\rho\mu(3\rho\mu\theta+4)}{\kappa^4(1+\rho\mu\theta)^2(2+\rho\mu\theta)^3}$ | $\frac{16\rho\mu(3\rho\mu\theta+4)}{\kappa^4(1+\rho\mu\theta)^2(2+\rho\mu\theta)^3}$ |
| See $\frac{\partial}{\partial\theta}c_{8,7}$ | See $\frac{\partial}{\partial\theta}c_{8,8}$ | See $\frac{\partial}{\partial\theta}c_{8,9}$ |

- $$\frac{\partial}{\partial\theta}c_{8,1} = -\frac{\begin{aligned}2(\kappa-1)\rho\mu\big\{(\rho\mu\theta)^2[-2(\rho\mu\theta)^3 - 3(\rho\mu\theta)^2 + 13(\rho\mu\theta) + 18] + \kappa(\rho\mu\theta)^2[2(\rho\mu\theta)^3 + 3(\rho\mu\theta)^2 - 13(\rho\mu\theta) - 18]\\ + \kappa^2[3(\rho\mu\theta)^4 + 13(\rho\mu\theta)^3 + 13(\rho\mu\theta)^2 - 3(\rho\mu\theta) - 6]\big\}\end{aligned}}{\kappa^4(1+\rho\mu\theta)^3(2+\rho\mu\theta)^3(3+\rho\mu\theta)^2}$$

- $\frac{\partial}{\partial\theta}c_{8,2} = \frac{\partial}{\partial\theta}c_{8,3} = \frac{\begin{array}{r}2\rho\mu\{(\rho\mu\theta)^2[-2(\rho\mu\theta)^3 - 3(\rho\mu\theta)^2 + 13(\rho\mu\theta) + 18] + \kappa(\rho\mu\theta)^2[2(\rho\mu\theta)^3 + 3(\rho\mu\theta)^2 - 13(\rho\mu\theta) - 18] \\ + \kappa^2[3(\rho\mu\theta)^4 + 13(\rho\mu\theta)^3 + 13(\rho\mu\theta)^2 - 3(\rho\mu\theta) - 6]\}\end{array}}{\kappa^4(1+\rho\mu\theta)^3(2+\rho\mu\theta)^3(3+\rho\mu\theta)^2}$
- $\frac{\partial}{\partial\theta}c_{8,4} = \frac{\begin{array}{l}\rho\mu\{2\rho\mu^2\theta^2[-2(\rho\mu\theta)^3 - 3(\rho\mu\theta)^2 + 13\rho\mu\theta + 18] \\ \quad + 2\kappa\mu\theta[2(\rho\mu\theta)^4 + 3(\rho+1)(\rho\mu\theta)^3 + (9\rho-13)(\rho\mu\theta)^2 - 2(\rho+9)(\rho\mu\theta) - 12\rho] \\ \quad + \kappa^2\rho^{-1}[-2(\rho\mu\theta)^5 - 3(3\rho+1)(\rho\mu\theta)^4 + (13-35\rho)(\rho\mu\theta)^3 - 6(4\rho-3)(\rho\mu\theta)^2 + 18\rho^2\mu\theta + 12\rho]\}\end{array}}{\kappa^4(1+\rho\mu\theta)^3(2+\rho\mu\theta)^3(3+\rho\mu\theta)^2}$
- $\frac{\partial}{\partial\theta}c_{8,5} = -\frac{2\rho\mu\{(\rho\mu\theta)^2[2(\rho\mu\theta)^3+3(\rho\mu\theta)^2-13\rho\mu\theta-18]+2\kappa\rho\mu\theta[3(\rho\mu\theta)^3+9(\rho\mu\theta)^2-2\rho\mu\theta-12]+\kappa^2[4(\rho\mu\theta)^3+15(\rho\mu\theta)^2+9\rho\mu\theta-6]\}}{\kappa^4(1+\rho\mu\theta)^3(2+\rho\mu\theta)^3(3+\rho\mu\theta)^2}$
- $\frac{\partial}{\partial\theta}c_{8,6} = \frac{\partial}{\partial\theta}c_{8,7} = \frac{2\rho\mu^2\theta\{\kappa[-3(\rho\mu\theta)^3-9(\rho\mu\theta)^2+2(\rho\mu\theta)+12]+\rho\mu\theta[-2(\rho\mu\theta)^3-3(\rho\mu\theta)^2+13(\rho\mu\theta)+18]\}}{\kappa^4(1+\rho\mu\theta)^3(2+\rho\mu\theta)^3(3+\rho\mu\theta)^2}$
- $\frac{\partial}{\partial\theta}c_{8,8} = \frac{2\rho\mu^2\theta\{\rho\mu\theta[-2(\rho\mu\theta)^3-3(\rho\mu\theta)^2+13\rho\mu\theta+18]+\kappa[(\rho\mu\theta)^4+3(\rho\mu\theta)^3-2(\rho\mu\theta)^2-10\rho\mu\theta-6]\}}{\kappa^4(1+\rho\mu\theta)^3(2+\rho\mu\theta)^3(3+\rho\mu\theta)^2}$
- $\frac{\partial}{\partial\theta}c_{8,9} = \frac{2\rho\mu^3\theta^2\{-2(\rho\mu\theta)^3-3(\rho\mu\theta)^2+13\rho\mu\theta+18\}}{\kappa^4(1+\rho\mu\theta)^3(2+\rho\mu\theta)^3(3+\rho\mu\theta)^2}$

##### A.1.2 Second derivatives

We derive the second-order partial derivatives of the log likelihood

$$\begin{pmatrix} \frac{\partial^2}{\partial\tau_1^2}l(\boldsymbol{\tau},\theta) & \frac{\partial^2}{\partial\tau_1\partial\tau_2}l(\boldsymbol{\tau},\theta) & \frac{\partial^2}{\partial\tau_1\partial\tau_3}l(\boldsymbol{\tau},\theta) & \frac{\partial^2}{\partial\tau_1\partial\tilde{\theta}}l(\boldsymbol{\tau},\theta) \\ \frac{\partial^2}{\partial\tau_1\partial\tau_2}l(\boldsymbol{\tau},\theta) & \frac{\partial^2}{\partial\tau_2^2}l(\boldsymbol{\tau},\theta) & \frac{\partial^2}{\partial\tau_2\partial\tau_3}l(\boldsymbol{\tau},\theta) & \frac{\partial^2}{\partial\tau_2\partial\tilde{\theta}}l(\boldsymbol{\tau},\theta) \\ \frac{\partial^2}{\partial\tau_1\partial\tau_3}l(\boldsymbol{\tau},\theta) & \frac{\partial^2}{\partial\tau_2\partial\tau_3}l(\boldsymbol{\tau},\theta) & \frac{\partial^2}{\partial\tau_3^2}l(\boldsymbol{\tau},\theta) & \frac{\partial^2}{\partial\tau_3\partial\tilde{\theta}}l(\boldsymbol{\tau},\theta) \\ \frac{\partial^2}{\partial\tau_1\partial\tilde{\theta}}l(\boldsymbol{\tau},\theta) & \frac{\partial^2}{\partial\tau_2\partial\tilde{\theta}}l(\boldsymbol{\tau},\theta) & \frac{\partial^2}{\partial\tau_3\partial\tilde{\theta}}l(\boldsymbol{\tau},\theta) & \frac{\partial^2}{\partial\tilde{\theta}^2}l(\boldsymbol{\tau},\theta) \end{pmatrix},$$

where

$$\begin{aligned} \frac{\partial^2}{\partial\tau_i\partial\tau_j}l(\boldsymbol{\tau},\theta) &= \sum_{k=1}^{15}\left\{-\frac{n_k^Q}{p_k^2}\cdot\frac{\partial}{\partial\tau_i}p_k\cdot\frac{\partial}{\partial\tau_j}p_k + \frac{n_k^Q}{p_k}\cdot\frac{\partial^2}{\partial\tau_i\partial\tau_j}p_k\right\} \\ &= \left(-\boldsymbol{n}^Q \oslash \boldsymbol{p}^2\right)^T\left(\frac{\partial}{\partial\tau_i}\boldsymbol{p}\odot\frac{\partial}{\partial\tau_j}\boldsymbol{p}\right) + \left(\boldsymbol{n}^Q \oslash \boldsymbol{p}\right)^T\left(\frac{\partial^2}{\partial\tau_i\partial\tau_j}\boldsymbol{p}\right) \text{ for } i,j \in \{1,2,3\}, \\ \frac{\partial^2}{\partial\tau_i\partial\tilde{\theta}}l(\boldsymbol{\tau},\theta) &= \sum_{k=1}^{15}\left\{-\frac{n_k^Q}{p_k^2}\cdot\frac{\partial}{\partial\tau_i}p_k\cdot\frac{\partial}{\partial\tilde{\theta}}p_k + \frac{n_k^Q}{p_k}\cdot\frac{\partial^2}{\partial\tau_i\partial\tilde{\theta}}p_k\right\} \\ &= \left(-\boldsymbol{n}^Q \oslash \boldsymbol{p}^2\right)^T\left(\frac{\partial}{\partial\tau_i}\boldsymbol{p}\odot\frac{\partial}{\partial\tilde{\theta}}\boldsymbol{p}\right) + \left(\boldsymbol{n}^Q \oslash \boldsymbol{p}\right)^T\left(\frac{\partial^2}{\partial\tau_i\partial\tilde{\theta}}\boldsymbol{p}\right) \text{ for } i = 1,2,3, \text{ and} \\ \frac{\partial^2}{\partial\tilde{\theta}^2}l(\boldsymbol{\tau},\theta) &= \sum_{k=1}^{15}\left\{-\frac{n_k^Q}{p_k^2}\left(\frac{\partial}{\partial\tilde{\theta}}p_k\right)^2 + \frac{n_k^Q}{p_k}\cdot\frac{\partial^2}{\partial\tilde{\theta}^2}p_k\right\} = \left(-\boldsymbol{n}^Q \oslash \boldsymbol{p}^2\right)^T\left(\frac{\partial}{\partial\tilde{\theta}}\boldsymbol{p}\right)^2 + \left(\boldsymbol{n}^Q \oslash \boldsymbol{p}\right)^T\left(\frac{\partial^2}{\partial\tilde{\theta}^2}\boldsymbol{p}\right). \end{aligned}$$

The second-order partial derivatives of the site pattern probabilities are derived as follows.

**Second-order partial derivatives for $\tau_i$ and $\tau_j$**

$$\frac{\partial^2}{\partial\tau_i\partial\tau_j}\boldsymbol{p} = \boldsymbol{W}_s\boldsymbol{C}_s^T\left(\frac{\partial^2}{\partial\tau_i\partial\tau_j}\boldsymbol{\beta}_s\right) = \boldsymbol{W}_s\boldsymbol{C}_s^T\left(\gamma_{\tau_i}\odot\gamma_{\tau_j}\odot\boldsymbol{\beta}_s\right) \text{ for } i,j \in \{1,2,3\}.$$

where $\gamma_{\tau_i}$ have been derived in A.1.1.

**Second-order partial derivatives for $\tau_i$ and $\tilde{\theta}$**

$$\begin{aligned}\frac{\partial^2}{\partial\tau_i\partial\tilde{\theta}}\boldsymbol{p} &= \boldsymbol{W}_s\left[\boldsymbol{C}_s^T\left(2\frac{\partial^2}{\partial\tau_i\partial\theta}\boldsymbol{\beta}_s\right)+\left(2\frac{\partial}{\partial\theta}\boldsymbol{C}\right)^T\left(\frac{\partial}{\partial\tau_i}\boldsymbol{\beta}_s\right)\right]\\ &= \boldsymbol{W}_s\left\{2\boldsymbol{C}_s^T\left[\gamma_{\tau_i\theta}\odot\boldsymbol{\beta}_s\right]+\left(2\frac{\partial}{\partial\theta}\boldsymbol{C}\right)^T\left(\gamma_{\tau_i}\odot\boldsymbol{\beta}_s\right)\right\}\ \text{for } i=1,2,3,\end{aligned}$$

where $\gamma_{\tau_i\theta} = \left[\frac{\partial}{\partial\theta}\gamma_{\tau_i}+\gamma_{\tau_i}\odot\gamma_\theta\right]$ and then we can derive

- $\gamma_{\tau_1\theta} = \gamma_{\tau_2\theta} = \left(0,0,0,0,0,0,0,0,-\frac{2(\theta+2\tau_1+2\tau_2-4\tau_3)}{\theta^3}\right)^T$, and
- $\gamma_{\tau_3\theta} = \left(0,0,0,0,0,0,0,0,\frac{4[\theta+(2\tau_1+2\tau_2-4\tau_3)(1+\rho\mu\theta)]}{\theta^3}\right)^T$.

**Second-order partial derivative for $\tilde{\theta}$ only**

$$\begin{aligned}\frac{\partial^2}{\partial\tilde{\theta}^2}\boldsymbol{p} &= \boldsymbol{W}_s\left[\boldsymbol{C}_s^T\left(4\frac{\partial^2}{\partial\theta^2}\boldsymbol{\beta}_s\right)+2\left(2\frac{\partial}{\partial\theta}\boldsymbol{C}_s\right)^T\left(2\frac{\partial}{\partial\theta}\boldsymbol{\beta}_s\right)+\left(4\frac{\partial^2}{\partial\theta^2}\boldsymbol{C}_s\right)^T\boldsymbol{\beta}_s\right]\\ &= \boldsymbol{W}_s\left[4\boldsymbol{C}_s^T\left(\gamma_{\theta^2}\odot\boldsymbol{\beta}_s\right)+8\left(\frac{\partial}{\partial\theta}\boldsymbol{C}_s\right)^T\left(\gamma_\theta\odot\boldsymbol{\beta}_s\right)+4\left(\frac{\partial^2}{\partial\theta^2}\boldsymbol{C}_s\right)^T\boldsymbol{\beta}_s\right]\end{aligned}$$

where $\gamma_\theta$ has been derived in A.1.1, $\gamma_{\theta^2} = \left[\frac{\partial}{\partial\theta}\gamma_\theta+\gamma_\theta\odot\gamma_\theta\right] = \left(0,0,0,0,0,0,0,0,\frac{4(\tau_1+\tau_2-2\tau_3)(\theta+\tau_1+\tau_2-2\tau_3)}{\theta^4}\right)^T$, and $\left(\frac{\partial^2}{\partial\theta^2}\boldsymbol{C}_s\right) = \left(\frac{\partial^2}{\partial\theta^2}\boldsymbol{c}_{s1},\frac{\partial^2}{\partial\theta^2}\boldsymbol{c}_{s2},...,\frac{\partial^2}{\partial\theta^2}\boldsymbol{c}_{s9}\right)$ is shown in table 2. Some derivatives in table 2 are shown separately as they are too long for the table.

- $\frac{\partial^2}{\partial\theta^2}c_{8,1} = \frac{\begin{aligned}&4(\kappa-1)\rho^2\mu^2\{(\rho\mu\theta)[-3(\rho\mu\theta)^6-9(\rho\mu\theta)^5+43(\rho\mu\theta)^4+201(\rho\mu\theta)^3+212(\rho\mu\theta)^2-36(\rho\mu\theta)-108]\\&+\kappa(\rho\mu\theta)[3(\rho\mu\theta)^6+9(\rho\mu\theta)^5-43(\rho\mu\theta)^4-201(\rho\mu\theta)^3-212(\rho\mu\theta)^2+36(\rho\mu\theta)+108]\\&+\kappa^2[6(\rho\mu\theta)^6+46(\rho\mu\theta)^5+117(\rho\mu\theta)^4+77(\rho\mu\theta)^3-123(\rho\mu\theta)^2-207(\rho\mu\theta)-84]\}\end{aligned}}{\kappa^4(1+\rho\mu\theta)^4(2+\rho\mu\theta)^4(3+\rho\mu\theta)^3}$
- $\frac{\partial^2}{\partial\theta^2}c_{8,2} = \frac{\partial}{\partial\theta}c_{8,3} = -\frac{\begin{aligned}&4\rho^2\mu^2\{(\rho\mu\theta)[-3(\rho\mu\theta)^6-9(\rho\mu\theta)^5+43(\rho\mu\theta)^4+201(\rho\mu\theta)^3+212(\rho\mu\theta)^2-36(\rho\mu\theta)-108]\\&+\kappa(\rho\mu\theta)[3(\rho\mu\theta)^6+9(\rho\mu\theta)^5-43(\rho\mu\theta)^4-201(\rho\mu\theta)^3-212(\rho\mu\theta)^2+36(\rho\mu\theta)+108]\\&+\kappa^2[6(\rho\mu\theta)^6+46(\rho\mu\theta)^5+117(\rho\mu\theta)^4+77(\rho\mu\theta)^3-123(\rho\mu\theta)^2-207(\rho\mu\theta)-84]\}\end{aligned}}{\kappa^4(1+\rho\mu\theta)^4(2+\rho\mu\theta)^4(3+\rho\mu\theta)^3}$
- $\frac{\partial^2}{\partial\theta^2}c_{8,4} = \frac{\begin{aligned}&2\rho^2\mu^2\{2\mu\theta[3(\rho\mu\theta)^6+9(\rho\mu\theta)^5-43(\rho\mu\theta)^4-201(\rho\mu\theta)^3-212(\rho\mu\theta)^2+36(\rho\mu\theta)+108]\\&+\kappa^2\rho^{-1}[3(\rho\mu\theta)^7+9(2\rho+1)(\rho\mu\theta)^6+(128\rho-43)(\rho\mu\theta)^5+3(92\rho-67)(\rho\mu\theta)^4\\&+2(23\rho-106)(\rho\mu\theta)^3-6(83\rho-6)(\rho\mu\theta)^2-18(29\rho-6)(\rho\mu\theta)-132\rho]\\&-2\kappa\rho^{-1}[3(\rho\mu\theta)^7+3(2\rho+3)(\rho\mu\theta)^6+(36\rho-43)(\rho\mu\theta)^5+3(14\rho-67)(\rho\mu\theta)^4\\&-4(27\rho+53)(\rho\mu\theta)^3-36(7\rho-1)(\rho\mu\theta)^2-108(\rho-1)(\rho\mu\theta)+36\rho]\}\end{aligned}}{\kappa^4(1+\rho\mu\theta)^3(2+\rho\mu\theta)^3(3+\rho\mu\theta)^2}$
- $\frac{\partial^2}{\partial\theta^2}c_{8,5} = \frac{\begin{aligned}&4\rho^2\mu^2\{\kappa^2[10(\rho\mu\theta)^5+75(\rho\mu\theta)^4+185(\rho\mu\theta)^3+129(\rho\mu\theta)^2-99(\rho\mu\theta)-120]\\&+12\kappa[(\rho\mu\theta)^6+6(\rho\mu\theta)^5+7(\rho\mu\theta)^4-18(\rho\mu\theta)^3-42(\rho\mu\theta)^2-18(\rho\mu\theta)+6]\\&+\rho\mu\theta[3(\rho\mu\theta)^6+9(\rho\mu\theta)^5-43(\rho\mu\theta)^4-201(\rho\mu\theta)^3-212(\rho\mu\theta)^2+36(\rho\mu\theta)+108]\}\end{aligned}}{\kappa^4(1+\rho\mu\theta)^3(2+\rho\mu\theta)^3(3+\rho\mu\theta)^2}$
- $\frac{\partial^2}{\partial\theta^2}c_{8,6} = \frac{\partial}{\partial\theta}c_{8,7} = \frac{\begin{aligned}&4\rho\mu^2\{6\kappa[(\rho\mu\theta)^6+6(\rho\mu\theta)^5+7(\rho\mu\theta)^4-18(\rho\mu\theta)^3-42(\rho\mu\theta)^2-18(\rho\mu\theta)+6]\\&+\rho\mu\theta[3(\rho\mu\theta)^6+9(\rho\mu\theta)^5-43(\rho\mu\theta)^4-201(\rho\mu\theta)^3-212(\rho\mu\theta)^2+36(\rho\mu\theta)+108]\}\end{aligned}}{\kappa^4(1+\rho\mu\theta)^3(2+\rho\mu\theta)^3(3+\rho\mu\theta)^2}$
- $\frac{\partial^2}{\partial\theta^2}c_{8,8} = -\frac{\begin{aligned}&2\rho\mu^2\{\kappa[3(\rho\mu\theta)^7+15(\rho\mu\theta)^6-7(\rho\mu\theta)^5-159(\rho\mu\theta)^4-320(\rho\mu\theta)^3-216(\rho\mu\theta)^2+36]\\&-2\rho\mu\theta[3(\rho\mu\theta)^6+9(\rho\mu\theta)^5-43(\rho\mu\theta)^4-201(\rho\mu\theta)^3-212(\rho\mu\theta)^2+36(\rho\mu\theta)+108]\}\end{aligned}}{\kappa^4(1+\rho\mu\theta)^3(2+\rho\mu\theta)^3(3+\rho\mu\theta)^2}$
- $\frac{\partial^2}{\partial\theta^2}c_{8,9} = \frac{4\rho\mu^3\theta[3(\rho\mu\theta)^6+9(\rho\mu\theta)^5-43(\rho\mu\theta)^4-201(\rho\mu\theta)^3-212(\rho\mu\theta)^2+36(\rho\mu\theta)+108]}{\kappa^4(1+\rho\mu\theta)^3(2+\rho\mu\theta)^3(3+\rho\mu\theta)^2}$

Table 2: Second-order partial derivatives of $\theta$ to the symmetric quartet coefficient matrix $\boldsymbol{C}$

| $\frac{\partial^2}{\partial\theta^2}\boldsymbol{c}_{s1}$ | $\frac{\partial^2}{\partial\theta^2}\boldsymbol{c}_{s2}$ | $\frac{\partial^2}{\partial\theta^2}\boldsymbol{c}_{s3}$ |
|---|---|---|
| $0$ | $0$ | $0$ |
| $\frac{2(\kappa-1)\rho^2\mu^2}{\kappa^4(1+\rho\mu\theta)^3}$ | $-\frac{2\rho^2\mu^2}{\kappa^4(1+\rho\mu\theta)^3}$ | $\frac{2(\kappa-1)\rho^2\mu^2}{\kappa^4(1+\rho\mu\theta)^3}$ |
| $\frac{2(\kappa-1)\rho^2\mu^2}{\kappa^4(1+\rho\mu\theta)^3}$ | $\frac{2(\kappa-1)\rho^2\mu^2}{\kappa^4(1+\rho\mu\theta)^3}$ | $-\frac{2\rho^2\mu^2}{\kappa^4(1+\rho\mu\theta)^3}$ |
| $\frac{6(\kappa-1)^2\rho^2\mu^2}{\kappa^4(1+\rho\mu\theta)^4}$ | $-\frac{6(\kappa-1)\rho^2\mu^2}{\kappa^4(1+\rho\mu\theta)^4}$ | $-\frac{6(\kappa-1)\rho^2\mu^2}{\kappa^4(1+\rho\mu\theta)^4}$ |
| $\frac{8(\kappa-1)\rho^2\mu^2}{\kappa^4(1+\rho\mu\theta)^3}$ | $\frac{4(\kappa-2)\rho^2\mu^2}{\kappa^4(1+\rho\mu\theta)^3}$ | $\frac{4(\kappa-2)\rho^2\mu^2}{\kappa^4(1+\rho\mu\theta)^3}$ |
| $\frac{8(\kappa-2)(\kappa-1)\rho^2\mu^2(3\rho^2\mu^2\theta^2+9\rho\mu\theta+7)}{\kappa^4(1+\rho\mu\theta)^3(2+\rho\mu\theta)^3}$ | $-\frac{8(\kappa-2)\rho^2\mu^2(3\rho^2\mu^2\theta^2+9\rho\mu\theta+7)}{\kappa^4(1+\rho\mu\theta)^3(2+\rho\mu\theta)^3}$ | $\frac{4(\kappa-2)^2\rho^2\mu^2(3\rho^2\mu^2\theta^2+9\rho\mu\theta+7)}{\kappa^4(1+\rho\mu\theta)^3(2+\rho\mu\theta)^3}$ |
| $\frac{8(\kappa-2)(\kappa-1)\rho^2\mu^2(3\rho^2\mu^2\theta^2+9\rho\mu\theta+7)}{\kappa^4(1+\rho\mu\theta)^3(2+\rho\mu\theta)^3}$ | $\frac{4(\kappa-2)^2\rho^2\mu^2(3\rho^2\mu^2\theta^2+9\rho\mu\theta+7)}{\kappa^4(1+\rho\mu\theta)^3(2+\rho\mu\theta)^3}$ | $-\frac{8(\kappa-2)\rho^2\mu^2(3\rho^2\mu^2\theta^2+9\rho\mu\theta+7)}{\kappa^4(1+\rho\mu\theta)^3(2+\rho\mu\theta)^3}$ |
| $\frac{8(\kappa-2)^2(\kappa-1)\rho^2\mu^2(6\rho^2\mu^2\theta^2+16\rho\mu\theta+11)}{\kappa^4(1+\rho\mu\theta)^3(2+\rho\mu\theta)^4}$ | $-\frac{8(\kappa-2)^2\rho^2\mu^2(6\rho^2\mu^2\theta^2+16\rho\mu\theta+11)}{\kappa^4(1+\rho\mu\theta)^3(2+\rho\mu\theta)^4}$ | $-\frac{8(\kappa-2)^2\rho^2\mu^2(6\rho^2\mu^2\theta^2+16\rho\mu\theta+11)}{\kappa^4(1+\rho\mu\theta)^3(2+\rho\mu\theta)^4}$ |
| See $\frac{\partial^2}{\partial\theta^2}c_{8,1}$ | See $\frac{\partial^2}{\partial\theta^2}c_{8,2}$ | See $\frac{\partial^2}{\partial\theta^2}c_{8,3}$ |

| $\frac{\partial^2}{\partial\theta^2}\boldsymbol{c}_{s4}$ | $\frac{\partial^2}{\partial\theta^2}\boldsymbol{c}_{s5}$ | $\frac{\partial^2}{\partial\theta^2}\boldsymbol{c}_{s6}$ |
|---|---|---|
| $0$ | $0$ | $0$ |
| $-\frac{2\rho^2\mu^2}{\kappa^4(1+\rho\mu\theta)^3}$ | $\frac{2(\kappa-1)\rho^2\mu^2}{\kappa^4(1+\rho\mu\theta)^3}$ | $-\frac{2\rho^2\mu^2}{\kappa^4(1+\rho\mu\theta)^3}$ |
| $-\frac{2\rho^2\mu^2}{\kappa^4(1+\rho\mu\theta)^3}$ | $\frac{2(\kappa-1)\rho^2\mu^2}{\kappa^4(1+\rho\mu\theta)^3}$ | $\frac{2(\kappa-1)\rho^2\mu^2}{\kappa^4(1+\rho\mu\theta)^3}$ |
| $\frac{6\rho^2\mu^2}{\kappa^4(1+\rho\mu\theta)^4}$ | $\frac{6(\kappa-1)^2\rho^2\mu^2}{\kappa^4(1+\rho\mu\theta)^4}$ | $-\frac{6(\kappa-1)\rho^2\mu^2}{\kappa^4(1+\rho\mu\theta)^4}$ |
| $\frac{4(\kappa-2)\rho^2\mu^2}{\kappa^4(1+\rho\mu\theta)^3}$ | $-\frac{8\rho^2\mu^2}{\kappa^4(1+\rho\mu\theta)^3}$ | $-\frac{8\rho^2\mu^2}{\kappa^4(1+\rho\mu\theta)^3}$ |
| $-\frac{8(\kappa-2)\rho^2\mu^2(3\rho^2\mu^2\theta^2+9\rho\mu\theta+7)}{\kappa^4(1+\rho\mu\theta)^3(2+\rho\mu\theta)^3}$ | $-\frac{8(\kappa-2)\rho^2\mu^2(3\rho^2\mu^2\theta^2+9\rho\mu\theta+7)}{\kappa^4(1+\rho\mu\theta)^3(2+\rho\mu\theta)^3}$ | $\frac{16\rho^2\mu^2(3\rho^2\mu^2\theta^2+9\rho\mu\theta+7)}{\kappa^4(1+\rho\mu\theta)^3(2+\rho\mu\theta)^3}$ |
| $-\frac{8(\kappa-2)\rho^2\mu^2(3\rho^2\mu^2\theta^2+9\rho\mu\theta+7)}{\kappa^4(1+\rho\mu\theta)^3(2+\rho\mu\theta)^3}$ | $-\frac{8(\kappa-2)\rho^2\mu^2(3\rho^2\mu^2\theta^2+9\rho\mu\theta+7)}{\kappa^4(1+\rho\mu\theta)^3(2+\rho\mu\theta)^3}$ | $-\frac{8(\kappa-2)\rho^2\mu^2(3\rho^2\mu^2\theta^2+9\rho\mu\theta+7)}{\kappa^4(1+\rho\mu\theta)^3(2+\rho\mu\theta)^3}$ |
| $\frac{16(\kappa-2)\rho^2\mu^2(6\rho^2\mu^2\theta^2+16\rho\mu\theta+11)}{\kappa^4(1+\rho\mu\theta)^3(2+\rho\mu\theta)^4}$ | $-\frac{8(\kappa-2)^2\rho^2\mu^2(6\rho^2\mu^2\theta^2+16\rho\mu\theta+11)}{\kappa^4(1+\rho\mu\theta)^3(2+\rho\mu\theta)^4}$ | $\frac{16(\kappa-2)\rho^2\mu^2(6\rho^2\mu^2\theta^2+16\rho\mu\theta+11)}{\kappa^4(1+\rho\mu\theta)^3(2+\rho\mu\theta)^4}$ |
| See $\frac{\partial^2}{\partial\theta^2}c_{8,4}$ | See $\frac{\partial^2}{\partial\theta^2}c_{8,5}$ | See $\frac{\partial^2}{\partial\theta^2}c_{8,6}$ |

| $\frac{\partial^2}{\partial\theta^2}\boldsymbol{c}_{s7}$ | $\frac{\partial^2}{\partial\theta^2}\boldsymbol{c}_{s8}$ | $\frac{\partial^2}{\partial\theta^2}\boldsymbol{c}_{s9}$ |
|---|---|---|
| $0$ | $0$ | $0$ |
| $\frac{2(\kappa-1)\rho^2\mu^2}{\kappa^4(1+\rho\mu\theta)^3}$ | $-\frac{2\rho^2\mu^2}{\kappa^4(1+\rho\mu\theta)^3}$ | $-\frac{2\rho^2\mu^2}{\kappa^4(1+\rho\mu\theta)^3}$ |
| $-\frac{2\rho^2\mu^2}{\kappa^4(1+\rho\mu\theta)^3}$ | $-\frac{2\rho^2\mu^2}{\kappa^4(1+\rho\mu\theta)^3}$ | $-\frac{2\rho^2\mu^2}{\kappa^4(1+\rho\mu\theta)^3}$ |
| $-\frac{6(\kappa-1)\rho^2\mu^2}{\kappa^4(1+\rho\mu\theta)^4}$ | $\frac{6\rho^2\mu^2}{\kappa^4(1+\rho\mu\theta)^4}$ | $\frac{6\rho^2\mu^2}{\kappa^4(1+\rho\mu\theta)^4}$ |
| $-\frac{8\rho^2\mu^2}{\kappa^4(1+\rho\mu\theta)^3}$ | $\frac{2(\kappa-4)\rho^2\mu^2}{\kappa^4(1+\rho\mu\theta)^3}$ | $-\frac{8\rho^2\mu^2}{\kappa^4(1+\rho\mu\theta)^3}$ |
| $-\frac{8(\kappa-2)\rho^2\mu^2(3\rho^2\mu^2\theta^2+9\rho\mu\theta+7)}{\kappa^4(1+\rho\mu\theta)^3(2+\rho\mu\theta)^3}$ | $-\frac{4(\kappa-4)\rho^2\mu^2(3\rho^2\mu^2\theta^2+9\rho\mu\theta+7)}{\kappa^4(1+\rho\mu\theta)^3(2+\rho\mu\theta)^3}$ | $\frac{16\rho^2\mu^2(3\rho^2\mu^2\theta^2+9\rho\mu\theta+7)}{\kappa^4(1+\rho\mu\theta)^3(2+\rho\mu\theta)^3}$ |
| $\frac{16\rho^2\mu^2(3\rho^2\mu^2\theta^2+9\rho\mu\theta+7)}{\kappa^4(1+\rho\mu\theta)^3(2+\rho\mu\theta)^3}$ | $-\frac{4(\kappa-4)\rho^2\mu^2(3\rho^2\mu^2\theta^2+9\rho\mu\theta+7)}{\kappa^4(1+\rho\mu\theta)^3(2+\rho\mu\theta)^3}$ | $\frac{16\rho^2\mu^2(3\rho^2\mu^2\theta^2+9\rho\mu\theta+7)}{\kappa^4(1+\rho\mu\theta)^3(2+\rho\mu\theta)^3}$ |
| $\frac{16(\kappa-2)\rho^2\mu^2(6\rho^2\mu^2\theta^2+16\rho\mu\theta+11)}{\kappa^4(1+\rho\mu\theta)^3(2+\rho\mu\theta)^4}$ | $\frac{8(\kappa-4)\rho^2\mu^2(6\rho^2\mu^2\theta^2+16\rho\mu\theta+11)}{\kappa^4(1+\rho\mu\theta)^3(2+\rho\mu\theta)^4}$ | $\frac{32\rho^2\mu^2(6\rho^2\mu^2\theta^2+16\rho\mu\theta+11)}{\kappa^4(1+\rho\mu\theta)^3(2+\rho\mu\theta)^4}$ |
| See $\frac{\partial^2}{\partial\theta^2}c_{8,7}$ | See $\frac{\partial^2}{\partial\theta^2}c_{8,8}$ | See $\frac{\partial^2}{\partial\theta^2}c_{8,9}$ |

#### A.2 Asymmetric quartet

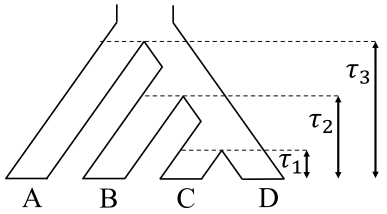


Figure 2: Asymmetric quartet according to Chifman and Kubatko (2015)

For an asymmetric quartet, the 15 category site pattern probabilities can be grouped into 11 bins:

| 1 | 2 | 3 | 4 | 5 | 6 | 7 | 8 | 9 | 10 | 11 |
|---|---|---|---|---|---|---|---|---|---|---|
| $xxxx$ | $xxxy$ | $xyxx$ | $yxxx$ | $xxyy$ | $xyxy$ | $xxyz$ | $yzxx$ | $xyxz$ | $yxxz$ | $xyzw$ |
| | $xxyx$ | | | | $yxxy$ | | | $xyzx$ | $yxzx$ | |

Similarly to the settings in A.1, the vector of 15 category site pattern probabilities can be written as

$$\boldsymbol{p} \equiv \left(p^{\rho}_{1|(Q_a,\boldsymbol{\tau},\theta)}, p^{\rho}_{2|(Q_a,\boldsymbol{\tau},\theta)}, ..., p^{\rho}_{15|(Q_a,\boldsymbol{\tau},\theta)}\right)^T = \boldsymbol{W}_a \boldsymbol{C}_a^T \boldsymbol{\beta}_a = \boldsymbol{W}_a (\boldsymbol{c}_{a1}^T \boldsymbol{\beta}_a, \boldsymbol{c}_{a2}^T \boldsymbol{\beta}_a, ..., \boldsymbol{c}_{a11}^T \boldsymbol{\beta}_a)^T,$$

where 1) $\boldsymbol{W}_a$ is a transformation weight matrix

$$\boldsymbol{W}_a = \begin{pmatrix} 4 & 0 & 0 & 0 & 0 & 0 & 0 & 0 & 0 & 0 & 0 \\ 0 & 12 & 0 & 0 & 0 & 0 & 0 & 0 & 0 & 0 & 0 \\ 0 & 12 & 0 & 0 & 0 & 0 & 0 & 0 & 0 & 0 & 0 \\ 0 & 0 & 12 & 0 & 0 & 0 & 0 & 0 & 0 & 0 & 0 \\ 0 & 0 & 0 & 12 & 0 & 0 & 0 & 0 & 0 & 0 & 0 \\ 0 & 0 & 0 & 0 & 0 & 12 & 0 & 0 & 0 & 0 & 0 \\ 0 & 0 & 0 & 0 & 0 & 12 & 0 & 0 & 0 & 0 & 0 \\ 0 & 0 & 0 & 0 & 12 & 0 & 0 & 0 & 0 & 0 & 0 \\ 0 & 0 & 0 & 0 & 0 & 0 & 0 & 0 & 24 & 0 & 0 \\ 0 & 0 & 0 & 0 & 0 & 0 & 0 & 0 & 24 & 0 & 0 \\ 0 & 0 & 0 & 0 & 0 & 0 & 0 & 0 & 0 & 24 & 0 \\ 0 & 0 & 0 & 0 & 0 & 0 & 0 & 0 & 0 & 24 & 0 \\ 0 & 0 & 0 & 0 & 0 & 0 & 24 & 0 & 0 & 0 & 0 \\ 0 & 0 & 0 & 0 & 0 & 0 & 0 & 24 & 0 & 0 & 0 \\ 0 & 0 & 0 & 0 & 0 & 0 & 0 & 0 & 0 & 0 & 24 \end{pmatrix}.$$

2) $\boldsymbol{C}_a = (\boldsymbol{c}_{a1}, \boldsymbol{c}_{a2}, ..., \boldsymbol{c}_{a11})$ be a 10×11 coefficient matrix, 3) $\boldsymbol{\beta}_a = \left(1, x_1^{2\rho\mu}, x_2^{2\rho\mu}, x_1^{\rho\mu}x_2^{2\rho\mu}, x_3^{2\rho\mu}, x_1^{\rho\mu}x_3^{2\rho\mu}, x_1^{2\rho\mu}x_3^{2\rho\mu}, x_2^{\rho\mu}x_3^{2\rho\mu}, x_1^{\rho\mu}x_2^{\rho\mu}x_3^{2\rho\mu}, x_1^{-\frac{2}{\theta}} x_2^{2(\rho\mu+\frac{1}{\theta})} x_3^{2\rho\mu}\right)^T$ be the $10 \times 1$ parameter vector, 4) $x_i = e^{-\tau_i}$ for $i \in \{1, 2, 3\}$ and 5) $\rho$ is the various evolution rate.

##### A.2.1 First derivatives

Referring to the equations of the first-order partial derivatives of the log likelihood in A.1.1, we derive the first-order partial derivatives of the site pattern probabilities as follows.

**First-order partial derivatives for $\tau_i$**

$$\frac{\partial}{\partial \tau_i}\boldsymbol{p} = \boldsymbol{W}_a \boldsymbol{C}_a^T \left(\gamma_{\tau_i} \odot \boldsymbol{\beta}_a\right) \text{ for } i = 1, 2, 3,$$

where

- $\gamma_{\tau_1} = \left(0, -2\rho\mu, 0, -\rho\mu, 0, -\rho\mu, -2\rho\mu, 0, -\rho\mu, \frac{2}{\theta}\right)^T$,
- $\gamma_{\tau_2} = \left(0, 0, -2\rho\mu, -2\rho\mu, 0, 0, 0, -\rho\mu, -\rho\mu, -2(\rho\mu + \frac{1}{\theta})\right)^T$, and
- $\gamma_{\tau_3} = \left(0, 0, 0, 0, -2\rho\mu, -2\rho\mu, -2\rho\mu, -2\rho\mu, -2\rho\mu, -2\rho\mu\right)^T$.

**First-order partial derivative for $\tilde{\theta}$**

$$\frac{\partial}{\partial\tilde{\theta}}\boldsymbol{p} = \boldsymbol{W}_a\left[2\boldsymbol{C}_a^T\left(\gamma_\theta \odot \boldsymbol{\beta}_a\right) + \left(2\frac{\partial}{\partial\theta}\boldsymbol{C}_a\right)^T \boldsymbol{\beta}_a\right]$$

where $\gamma_\theta = \left(0, 0, 0, 0, 0, 0, 0, 0, 0, \frac{2(\tau_2 - \tau_1)}{\theta^2}\right)^T$, and $\left(\frac{\partial}{\partial\theta}\boldsymbol{C}_a\right) = \left(\frac{\partial}{\partial\theta}\boldsymbol{c}_{a1}, \frac{\partial}{\partial\theta}\boldsymbol{c}_{a2}, ..., \frac{\partial}{\partial\theta}\boldsymbol{c}_{a11}\right)$ is shown in table 3. Some derivatives in table 3 are shown separately as they are too long for the table.

- $\frac{\partial}{\partial\theta}c_{9,1} = -\frac{2(\kappa-1)\rho\mu\{(\rho\mu\theta)^2[-2(\rho\mu\theta)^3 - 3(\rho\mu\theta)^2 + 13(\rho\mu\theta) + 18] + \kappa(\rho\mu\theta)^2[2(\rho\mu\theta)^3 + 3(\rho\mu\theta)^2 - 13(\rho\mu\theta) - 18] + \kappa^2[3(\rho\mu\theta)^4 + 13(\rho\mu\theta)^3 + 13(\rho\mu\theta)^2 - 3(\rho\mu\theta) - 6]\}}{\kappa^4(1+\rho\mu\theta)^3(2+\rho\mu\theta)^3(3+\rho\mu\theta)^2}$
- $\frac{\partial}{\partial\theta}c_{9,2} = \frac{\partial}{\partial\theta}c_{9,3} = \frac{\partial}{\partial\theta}c_{9,4} = \frac{2\rho\mu\{(\rho\mu\theta)^2[-2(\rho\mu\theta)^3 - 3(\rho\mu\theta)^2 + 13(\rho\mu\theta) + 18] + \kappa(\rho\mu\theta)^2[2(\rho\mu\theta)^3 + 3(\rho\mu\theta)^2 - 13(\rho\mu\theta) - 18] + \kappa^2[3(\rho\mu\theta)^4 + 13(\rho\mu\theta)^3 + 13(\rho\mu\theta)^2 - 3(\rho\mu\theta) - 6]\}}{\kappa^4(1+\rho\mu\theta)^3(2+\rho\mu\theta)^3(3+\rho\mu\theta)^2}$
- $\frac{\partial}{\partial\theta}c_{9,5} = -\frac{2\rho\mu\{(\rho\mu\theta)^2[2(\rho\mu\theta)^3+3(\rho\mu\theta)^2-13\rho\mu\theta-18]+2\kappa\rho\mu\theta[3(\rho\mu\theta)^3+9(\rho\mu\theta)^2-2\rho\mu\theta-12]+\kappa^2[4(\rho\mu\theta)^3+15(\rho\mu\theta)^2+9\rho\mu\theta-6]\}}{\kappa^4(1+\rho\mu\theta)^3(2+\rho\mu\theta)^3(3+\rho\mu\theta)^2}$
- $\frac{\partial}{\partial\theta}c_{9,6} = \frac{\rho\mu\{2\rho\mu^2\theta^2[-2(\rho\mu\theta)^3 - 3(\rho\mu\theta)^2 + 13\rho\mu\theta + 18] + 2\kappa\mu\theta[2(\rho\mu\theta)^4 + 3(\rho+1)(\rho\mu\theta)^3 + (9\rho-13)(\rho\mu\theta)^2 - 2(\rho+9)(\rho\mu\theta) - 12\rho] + \kappa^2\rho^{-1}[-2(\rho\mu\theta)^5 - 3(3\rho+1)(\rho\mu\theta)^4 + (13-35\rho)(\rho\mu\theta)^3 - 6(4\rho-3)(\rho\mu\theta)^2 + 18\rho^2\mu\theta + 12\rho]\}}{\kappa^4(1+\rho\mu\theta)^3(2+\rho\mu\theta)^3(3+\rho\mu\theta)^2}$
- $\frac{\partial}{\partial\theta}c_{9,7} = \frac{\partial}{\partial\theta}c_{9,8} = \frac{2\rho\mu^2\theta\{\kappa[-3(\rho\mu\theta)^3-9(\rho\mu\theta)^2+2(\rho\mu\theta)+12]+\rho\mu\theta[-2(\rho\mu\theta)^3-3(\rho\mu\theta)^2+13(\rho\mu\theta)+18]\}}{\kappa^4(1+\rho\mu\theta)^3(2+\rho\mu\theta)^3(3+\rho\mu\theta)^2}$
- $\frac{\partial}{\partial\theta}c_{9,9} = \frac{\partial}{\partial\theta}c_{9,10} = \frac{2\rho\mu^2\theta\{\rho\mu\theta[-2(\rho\mu\theta)^3-3(\rho\mu\theta)^2+13\rho\mu\theta+18]+\kappa[(\rho\mu\theta)^4+3(\rho\mu\theta)^3-2(\rho\mu\theta)^2-10\rho\mu\theta-6]\}}{\kappa^4(1+\rho\mu\theta)^3(2+\rho\mu\theta)^3(3+\rho\mu\theta)^2}$
- $\frac{\partial}{\partial\theta}c_{9,11} = \frac{2\rho\mu^3\theta^2\{-2(\rho\mu\theta)^3-3(\rho\mu\theta)^2+13\rho\mu\theta+18\}}{\kappa^4(1+\rho\mu\theta)^3(2+\rho\mu\theta)^3(3+\rho\mu\theta)^2}$

Table 3: First-order partial derivatives of $\theta$ to the asymmetric quartet coefficient matrix $\boldsymbol{C}_a$

| $\frac{\partial}{\partial\theta}\boldsymbol{c}_{a1}$ | $\frac{\partial}{\partial\theta}\boldsymbol{c}_{a2}$ | $\frac{\partial}{\partial\theta}\boldsymbol{c}_{a3}$ | $\frac{\partial}{\partial\theta}\boldsymbol{c}_{a4}$ |
|---|---|---|---|
| $0$ | $0$ | $0$ | $0$ |
| $-\frac{(\kappa-1)\rho\mu}{\kappa^4(1+\rho\mu\theta)^2}$ | $\frac{\rho\mu}{\kappa^4(1+\rho\mu\theta)^2}$ | $-\frac{(\kappa-1)\rho\mu}{\kappa^4(1+\rho\mu\theta)^2}$ | $-\frac{(\kappa-1)\rho\mu}{\kappa^4(1+\rho\mu\theta)^2}$ |
| $-\frac{2(\kappa-1)\rho\mu}{\kappa^4(1+\rho\mu\theta)^2}$ | $-\frac{(\kappa-2)\rho\mu}{\kappa^4(1+\rho\mu\theta)^2}$ | $\frac{2\rho\mu}{\kappa^4(1+\rho\mu\theta)^2}$ | $-\frac{2(\kappa-1)\rho\mu}{\kappa^4(1+\rho\mu\theta)^2}$ |
| $-\frac{2(\kappa-2)(\kappa-1)\rho\mu(2\rho\mu\theta+3)}{\kappa^4(1+\rho\mu\theta)^2(2+\rho\mu\theta)^2}$ | $\frac{2(\kappa-2)\rho\mu(2\rho\mu\theta+3)}{\kappa^4(1+\rho\mu\theta)^2(2+\rho\mu\theta)^2}$ | $\frac{2(\kappa-2)\rho\mu(2\rho\mu\theta+3)}{\kappa^4(1+\rho\mu\theta)^2(2+\rho\mu\theta)^2}$ | $-\frac{2(\kappa-2)(\kappa-1)\rho\mu(2\rho\mu\theta+3)}{\kappa^4(1+\rho\mu\theta)^2(2+\rho\mu\theta)^2}$ |
| $-\frac{3(\kappa-1)\rho\mu}{\kappa^4(1+\rho\mu\theta)^2}$ | $-\frac{(2\kappa-3)\rho\mu}{\kappa^4(1+\rho\mu\theta)^2}$ | $-\frac{(2\kappa-3)\rho\mu}{\kappa^4(1+\rho\mu\theta)^2}$ | $\frac{3\rho\mu}{\kappa^4(1+\rho\mu\theta)^2}$ |
| $-\frac{2(\kappa-2)(\kappa-1)\rho\mu(2\rho\mu\theta+3)}{\kappa^4(1+\rho\mu\theta)^2(2+\rho\mu\theta)^2}$ | $\frac{2(\kappa-2)\rho\mu(2\rho\mu\theta+3)}{\kappa^4(1+\rho\mu\theta)^2(2+\rho\mu\theta)^2}$ | $-\frac{2(\kappa-2)(\kappa-1)\rho\mu(2\rho\mu\theta+3)}{\kappa^4(1+\rho\mu\theta)^2(2+\rho\mu\theta)^2}$ | $\frac{2(\kappa-2)\rho\mu(2\rho\mu\theta+3)}{\kappa^4(1+\rho\mu\theta)^2(2+\rho\mu\theta)^2}$ |
| $-\frac{2(\kappa-1)^2\rho\mu}{\kappa^4(1+\rho\mu\theta)^3}$ | $\frac{2(\kappa-1)\rho\mu}{\kappa^4(1+\rho\mu\theta)^3}$ | $\frac{2(\kappa-1)\rho\mu}{\kappa^4(1+\rho\mu\theta)^3}$ | $\frac{2(\kappa-1)\rho\mu}{\kappa^4(1+\rho\mu\theta)^3}$ |
| $-\frac{4(\kappa-2)(\kappa-1)\rho\mu(2\rho\mu\theta+3)}{\kappa^4(1+\rho\mu\theta)^2(2+\rho\mu\theta)^2}$ | $-\frac{2(\kappa-2)^2\rho\mu(2\rho\mu\theta+3)}{\kappa^4(1+\rho\mu\theta)^2(2+\rho\mu\theta)^2}$ | $\frac{4(\kappa-2)\rho\mu(2\rho\mu\theta+3)}{\kappa^4(1+\rho\mu\theta)^2(2+\rho\mu\theta)^2}$ | $\frac{4(\kappa-2)\rho\mu(2\rho\mu\theta+3)}{\kappa^4(1+\rho\mu\theta)^2(2+\rho\mu\theta)^2}$ |
| $-\frac{4(\kappa-2)^2(\kappa-1)\rho\mu(3\rho\mu\theta+4)}{\kappa^4(1+\rho\mu\theta)^2(2+\rho\mu\theta)^3}$ | $\frac{4(\kappa-2)^2\rho\mu(3\rho\mu\theta+4)}{\kappa^4(1+\rho\mu\theta)^2(2+\rho\mu\theta)^3}$ | $\frac{4(\kappa-2)^2\rho\mu(3\rho\mu\theta+4)}{\kappa^4(1+\rho\mu\theta)^2(2+\rho\mu\theta)^3}$ | $\frac{4(\kappa-2)^2\rho\mu(3\rho\mu\theta+4)}{\kappa^4(1+\rho\mu\theta)^2(2+\rho\mu\theta)^3}$ |
| See $\frac{\partial}{\partial\theta}c_{9,1}$ | See $\frac{\partial}{\partial\theta}c_{9,2}$ | See $\frac{\partial}{\partial\theta}c_{9,3}$ | See $\frac{\partial}{\partial\theta}c_{9,4}$ |

| $\frac{\partial}{\partial\theta}\boldsymbol{c}_{a5}$ | $\frac{\partial}{\partial\theta}\boldsymbol{c}_{a6}$ | $\frac{\partial}{\partial\theta}\boldsymbol{c}_{a7}$ | $\frac{\partial}{\partial\theta}\boldsymbol{c}_{a8}$ |
|---|---|---|---|
| $0$ | $0$ | $0$ | $0$ |
| $-\frac{(\kappa-1)\rho\mu}{\kappa^4(1+\rho\mu\theta)^2}$ | $\frac{\rho\mu}{\kappa^4(1+\rho\mu\theta)^2}$ | $\frac{\rho\mu}{\kappa^4(1+\rho\mu\theta)^2}$ | $-\frac{(\kappa-1)\rho\mu}{\kappa^4(1+\rho\mu\theta)^2}$ |
| $\frac{2\rho\mu}{\kappa^4(1+\rho\mu\theta)^2}$ | $-\frac{(\kappa-2)\rho\mu}{\kappa^4(1+\rho\mu\theta)^2}$ | $\frac{2\rho\mu}{\kappa^4(1+\rho\mu\theta)^2}$ | $\frac{2\rho\mu}{\kappa^4(1+\rho\mu\theta)^2}$ |
| $\frac{2(\kappa-2)\rho\mu(2\rho\mu\theta+3)}{\kappa^4(1+\rho\mu\theta)^2(2+\rho\mu\theta)^2}$ | $\frac{2(\kappa-2)\rho\mu(2\rho\mu\theta+3)}{\kappa^4(1+\rho\mu\theta)^2(2+\rho\mu\theta)^2}$ | $-\frac{4\rho\mu(2\rho\mu\theta+3)}{\kappa^4(1+\rho\mu\theta)^2(2+\rho\mu\theta)^2}$ | $\frac{2(\kappa-2)\rho\mu(2\rho\mu\theta+3)}{\kappa^4(1+\rho\mu\theta)^2(2+\rho\mu\theta)^2}$ |
| $-\frac{(\kappa-3)\rho\mu}{\kappa^4(1+\rho\mu\theta)^2}$ | $-\frac{(\kappa-3)\rho\mu}{\kappa^4(1+\rho\mu\theta)^2}$ | $-\frac{(\kappa-3)\rho\mu}{\kappa^4(1+\rho\mu\theta)^2}$ | $\frac{3\rho\mu}{\kappa^4(1+\rho\mu\theta)^2}$ |
| $\frac{2(\kappa-2)\rho\mu(2\rho\mu\theta+3)}{\kappa^4(1+\rho\mu\theta)^2(2+\rho\mu\theta)^2}$ | $\frac{2(\kappa-2)\rho\mu(2\rho\mu\theta+3)}{\kappa^4(1+\rho\mu\theta)^2(2+\rho\mu\theta)^2}$ | $-\frac{4\rho\mu(2\rho\mu\theta+3)}{\kappa^4(1+\rho\mu\theta)^2(2+\rho\mu\theta)^2}$ | $\frac{2(\kappa-2)\rho\mu(2\rho\mu\theta+3)}{\kappa^4(1+\rho\mu\theta)^2(2+\rho\mu\theta)^2}$ |
| $-\frac{2(\kappa-1)^2\rho\mu}{\kappa^4(1+\rho\mu\theta)^3}$ | $-\frac{2\rho\mu}{\kappa^4(1+\rho\mu\theta)^3}$ | $\frac{2(\kappa-1)\rho\mu}{\kappa^4(1+\rho\mu\theta)^3}$ | $\frac{2(\kappa-1)\rho\mu}{\kappa^4(1+\rho\mu\theta)^3}$ |
| $\frac{4(\kappa-2)\rho\mu(2\rho\mu\theta+3)}{\kappa^4(1+\rho\mu\theta)^2(2+\rho\mu\theta)^2}$ | $\frac{4(\kappa-2)\rho\mu(2\rho\mu\theta+3)}{\kappa^4(1+\rho\mu\theta)^2(2+\rho\mu\theta)^2}$ | $\frac{4(\kappa-2)\rho\mu(2\rho\mu\theta+3)}{\kappa^4(1+\rho\mu\theta)^2(2+\rho\mu\theta)^2}$ | $-\frac{8\rho\mu(2\rho\mu\theta+3)}{\kappa^4(1+\rho\mu\theta)^2(2+\rho\mu\theta)^2}$ |
| $\frac{4(\kappa-2)^2\rho\mu(3\rho\mu\theta+4)}{\kappa^4(1+\rho\mu\theta)^2(2+\rho\mu\theta)^3}$ | $-\frac{8(\kappa-2)\rho\mu(3\rho\mu\theta+4)}{\kappa^4(1+\rho\mu\theta)^2(2+\rho\mu\theta)^3}$ | $-\frac{8(\kappa-2)\rho\mu(3\rho\mu\theta+4)}{\kappa^4(1+\rho\mu\theta)^2(2+\rho\mu\theta)^3}$ | $-\frac{8(\kappa-2)\rho\mu(3\rho\mu\theta+4)}{\kappa^4(1+\rho\mu\theta)^2(2+\rho\mu\theta)^3}$ |
| See $\frac{\partial}{\partial\theta}c_{9,5}$ | See $\frac{\partial}{\partial\theta}c_{9,6}$ | See $\frac{\partial}{\partial\theta}c_{9,7}$ | See $\frac{\partial}{\partial\theta}c_{9,8}$ |

| $\frac{\partial}{\partial\theta}\boldsymbol{c}_{a9}$ | $\frac{\partial}{\partial\theta}\boldsymbol{c}_{a10}$ | $\frac{\partial}{\partial\theta}\boldsymbol{c}_{a11}$ |
|---|---|---|
| $0$ | $0$ | $0$ |
| $\frac{\rho\mu}{\kappa^4(1+\rho\mu\theta)^2}$ | $\frac{\rho\mu}{\kappa^4(1+\rho\mu\theta)^2}$ | $\frac{\rho\mu}{\kappa^4(1+\rho\mu\theta)^2}$ |
| $\frac{2\rho\mu}{\kappa^4(1+\rho\mu\theta)^2}$ | $-\frac{(\kappa-2)\rho\mu}{\kappa^4(1+\rho\mu\theta)^2}$ | $\frac{2\rho\mu}{\kappa^4(1+\rho\mu\theta)^2}$ |
| $-\frac{4\rho\mu(2\rho\mu\theta+3)}{\kappa^4(1+\rho\mu\theta)^2(2+\rho\mu\theta)^2}$ | $\frac{2(\kappa-2)\rho\mu(2\rho\mu\theta+3)}{\kappa^4(1+\rho\mu\theta)^2(2+\rho\mu\theta)^2}$ | $-\frac{4\rho\mu(2\rho\mu\theta+3)}{\kappa^4(1+\rho\mu\theta)^2(2+\rho\mu\theta)^2}$ |
| $-\frac{(\kappa-3)\rho\mu}{\kappa^4(1+\rho\mu\theta)^2}$ | $\frac{3\rho\mu}{\kappa^4(1+\rho\mu\theta)^2}$ | $\frac{3\rho\mu}{\kappa^4(1+\rho\mu\theta)^2}$ |
| $\frac{2(\kappa-2)\rho\mu(2\rho\mu\theta+3)}{\kappa^4(1+\rho\mu\theta)^2(2+\rho\mu\theta)^2}$ | $-\frac{4\rho\mu(2\rho\mu\theta+3)}{\kappa^4(1+\rho\mu\theta)^2(2+\rho\mu\theta)^2}$ | $-\frac{4\rho\mu(2\rho\mu\theta+3)}{\kappa^4(1+\rho\mu\theta)^2(2+\rho\mu\theta)^2}$ |
| $-\frac{2\rho\mu}{\kappa^4(1+\rho\mu\theta)^3}$ | $-\frac{2\rho\mu}{\kappa^4(1+\rho\mu\theta)^3}$ | $-\frac{2\rho\mu}{\kappa^4(1+\rho\mu\theta)^3}$ |
| $\frac{2(\kappa-4)\rho\mu(2\rho\mu\theta+3)}{\kappa^4(1+\rho\mu\theta)^2(2+\rho\mu\theta)^2}$ | $\frac{2(\kappa-4)\rho\mu(2\rho\mu\theta+3)}{\kappa^4(1+\rho\mu\theta)^2(2+\rho\mu\theta)^2}$ | $-\frac{8\rho\mu(2\rho\mu\theta+3)}{\kappa^4(1+\rho\mu\theta)^2(2+\rho\mu\theta)^2}$ |
| $-\frac{4(\kappa-4)\rho\mu(3\rho\mu\theta+4)}{\kappa^4(1+\rho\mu\theta)^2(2+\rho\mu\theta)^3}$ | $-\frac{4(\kappa-4)\rho\mu(3\rho\mu\theta+4)}{\kappa^4(1+\rho\mu\theta)^2(2+\rho\mu\theta)^3}$ | $\frac{16\rho\mu(3\rho\mu\theta+4)}{\kappa^4(1+\rho\mu\theta)^2(2+\rho\mu\theta)^3}$ |
| See $\frac{\partial}{\partial\theta}c_{9,9}$ | See $\frac{\partial}{\partial\theta}c_{9,10}$ | See $\frac{\partial}{\partial\theta}c_{9,11}$ |

##### A.2.2 Second derivatives

Referring to the Hessian matrix that is the second-order partial derivatives of the log likelihood in A.1.2, we derive the second-order partial derivatives of the site pattern probabilities as follows.

**Second-order partial derivatives for $\tau_i$ and $\tau_j$**

$$\frac{\partial^2}{\partial\tau_i\partial\tau_j}\boldsymbol{p} = \boldsymbol{W}_a\boldsymbol{C}_a^T\left(\gamma_{\tau_i}\odot\gamma_{\tau_j}\odot\boldsymbol{\beta}_a\right) \text{ for } i,j\in\{1,2,3\}.$$

where $\gamma_{\tau_i}$ have been derived in A.2.1.

**Second-order partial derivatives for $\tau_i$ and $\tilde{\theta}$**

$$\frac{\partial^2}{\partial\tau_i\partial\tilde{\theta}}\boldsymbol{p} = \boldsymbol{W}_a\left\{2\boldsymbol{C}_a^T\left[\gamma_{\tau_i\theta}\odot\boldsymbol{\beta}_a\right] + \left(2\frac{\partial}{\partial\theta}\boldsymbol{C}_a\right)^T\left(\gamma_{\tau_i}\odot\boldsymbol{\beta}_a\right)\right\}, \text{ for } i=1,2,3,$$

where $\gamma_{\tau_i\theta} = \left[\frac{\partial}{\partial\theta}\gamma_{\tau_i} + \gamma_{\tau_i}\odot\gamma_\theta\right]$ and then we can derive

- $\gamma_{\tau_1\theta} = \left(0,0,0,0,0,0,0,0,0,-\frac{2(\theta+2\tau_1-2\tau_2)}{\theta^3}\right)^T$,
- $\gamma_{\tau_2\theta} = \left(0,0,0,0,0,0,0,0,0,\frac{2[\theta+2(1+\rho\mu\theta)(\tau_1-\tau_2)]}{\theta^3}\right)^T$, and
- $\gamma_{\tau_3\theta} = \left(0,0,0,0,0,0,0,0,0,\frac{4\rho\mu(\tau_1-\tau_2)}{\theta^2}\right)^T$.

**Second-order partial derivative for $\tilde{\theta}$ only**

$$\frac{\partial^2}{\partial\tilde{\theta}^2}\boldsymbol{p} = \boldsymbol{W}_a\left[4\boldsymbol{C}_a^T\left(\gamma_{\theta^2}\odot\boldsymbol{\beta}_a\right) + 8\left(\frac{\partial}{\partial\theta}\boldsymbol{C}_a\right)^T\left(\gamma_\theta\odot\boldsymbol{\beta}_a\right) + 4\left(\frac{\partial^2}{\partial\theta^2}\boldsymbol{C}_a\right)^T\boldsymbol{\beta}_a\right]$$

where $\gamma_\theta$ has been derived in A.2.1, $\gamma_{\theta^2} = \left(\frac{\partial}{\partial\theta}\gamma_\theta + \gamma_\theta^2\right) = \left(0,0,0,0,0,0,0,0,\frac{4(\tau_1-\tau_2)(\theta+\tau_1-\tau_2)}{\theta^4}\right)^T$ and $\left(\frac{\partial^2}{\partial\theta^2}\boldsymbol{C}_a\right) = \left(\frac{\partial^2}{\partial\theta^2}\boldsymbol{c}_{a1},\frac{\partial^2}{\partial\theta^2}\boldsymbol{c}_{a2},\ldots,\frac{\partial^2}{\partial\theta^2}\boldsymbol{c}_{a11}\right)$ is shown in the table. Some derivatives in $\left(\frac{\partial^2}{\partial\theta^2}\boldsymbol{C}_a\right)$ are shown separately as they are too long for the table.

- $\frac{\partial^2}{\partial\theta^2}c_{9,1} = \frac{4(\kappa-1)\rho^2\mu^2\{(\rho\mu\theta)[-3(\rho\mu\theta)^6-9(\rho\mu\theta)^5+43(\rho\mu\theta)^4+201(\rho\mu\theta)^3+212(\rho\mu\theta)^2-36(\rho\mu\theta)-108] + \kappa(\rho\mu\theta)[3(\rho\mu\theta)^6+9(\rho\mu\theta)^5-43(\rho\mu\theta)^4-201(\rho\mu\theta)^3-212(\rho\mu\theta)^2+36(\rho\mu\theta)+108] + \kappa^2[6(\rho\mu\theta)^6+46(\rho\mu\theta)^5+117(\rho\mu\theta)^4+77(\rho\mu\theta)^3-123(\rho\mu\theta)^2-207(\rho\mu\theta)-84]\}}{\kappa^4(1+\rho\mu\theta)^4(2+\rho\mu\theta)^4(3+\rho\mu\theta)^3}$
- $\frac{\partial^2}{\partial\theta^2}c_{9,2} = \frac{\partial}{\partial\theta}c_{9,3} = \frac{\partial}{\partial\theta}c_{9,4} = -\frac{4\rho^2\mu^2\{(\rho\mu\theta)[-3(\rho\mu\theta)^6-9(\rho\mu\theta)^5+43(\rho\mu\theta)^4+201(\rho\mu\theta)^3+212(\rho\mu\theta)^2-36(\rho\mu\theta)-108] + \kappa(\rho\mu\theta)[3(\rho\mu\theta)^6+9(\rho\mu\theta)^5-43(\rho\mu\theta)^4-201(\rho\mu\theta)^3-212(\rho\mu\theta)^2+36(\rho\mu\theta)+108] + \kappa^2[6(\rho\mu\theta)^6+46(\rho\mu\theta)^5+117(\rho\mu\theta)^4+77(\rho\mu\theta)^3-123(\rho\mu\theta)^2-207(\rho\mu\theta)-84]\}}{\kappa^4(1+\rho\mu\theta)^4(2+\rho\mu\theta)^4(3+\rho\mu\theta)^3}$
- $\frac{\partial^2}{\partial\theta^2}c_{9,5} = \frac{4\rho^2\mu^2\{\kappa^2[10(\rho\mu\theta)^5+75(\rho\mu\theta)^4+185(\rho\mu\theta)^3+129(\rho\mu\theta)^2-99(\rho\mu\theta)-120] + 12\kappa[(\rho\mu\theta)^6+6(\rho\mu\theta)^5+7(\rho\mu\theta)^4-18(\rho\mu\theta)^3-42(\rho\mu\theta)^2-18(\rho\mu\theta)+6] + \rho\mu\theta[3(\rho\mu\theta)^6+9(\rho\mu\theta)^5-43(\rho\mu\theta)^4-201(\rho\mu\theta)^3-212(\rho\mu\theta)^2+36(\rho\mu\theta)+108]\}}{\kappa^4(1+\rho\mu\theta)^4(2+\rho\mu\theta)^4(3+\rho\mu\theta)^3}$
- $\frac{\partial^2}{\partial\theta^2}c_{9,6} = \frac{2\rho^2\mu^2\{2\mu\theta[3(\rho\mu\theta)^6+9(\rho\mu\theta)^5-43(\rho\mu\theta)^4-201(\rho\mu\theta)^3-212(\rho\mu\theta)^2+36(\rho\mu\theta)+108] + \kappa^2\rho^{-1}[3(\rho\mu\theta)^7+9(2\rho+1)(\rho\mu\theta)^6+(128\rho-43)(\rho\mu\theta)^5+3(92\rho-67)(\rho\mu\theta)^4 + 2(23\rho-106)(\rho\mu\theta)^3-6(83\rho-6)(\rho\mu\theta)^2-18(29\rho-6)(\rho\mu\theta)-132\rho] - 2\kappa\rho^{-1}[3(\rho\mu\theta)^7+3(2\rho+3)(\rho\mu\theta)^6+(36\rho-43)(\rho\mu\theta)^5+3(14\rho-67)(\rho\mu\theta)^4 - 4(27\rho+53)(\rho\mu\theta)^3-36(7\rho-1)(\rho\mu\theta)^2-108(\rho-1)(\rho\mu\theta)+36\rho]\}}{\kappa^4(1+\rho\mu\theta)^4(2+\rho\mu\theta)^4(3+\rho\mu\theta)^3}$

- $\frac{\partial^2}{\partial\theta^2}c_{9,7} = \frac{\partial}{\partial\theta}c_{9,8} = \frac{4\rho\mu^2\{6\kappa[(\rho\mu\theta)^6+6(\rho\mu\theta)^5+7(\rho\mu\theta)^4-18(\rho\mu\theta)^3-42(\rho\mu\theta)^2-18(\rho\mu\theta)+6] + \rho\mu\theta[3(\rho\mu\theta)^6+9(\rho\mu\theta)^5-43(\rho\mu\theta)^4-201(\rho\mu\theta)^3-212(\rho\mu\theta)^2+36(\rho\mu\theta)+108]\}}{\kappa^4(1+\rho\mu\theta)^4(2+\rho\mu\theta)^4(3+\rho\mu\theta)^3}$
- $\frac{\partial^2}{\partial\theta^2}c_{9,9} = \frac{\partial^2}{\partial\theta^2}c_{9,10} = -\frac{2\rho\mu^2\{\kappa[3(\rho\mu\theta)^7+15(\rho\mu\theta)^6-7(\rho\mu\theta)^5-159(\rho\mu\theta)^4-320(\rho\mu\theta)^3-216(\rho\mu\theta)^2+36] - 2\rho\mu\theta[3(\rho\mu\theta)^6+9(\rho\mu\theta)^5-43(\rho\mu\theta)^4-201(\rho\mu\theta)^3-212(\rho\mu\theta)^2+36(\rho\mu\theta)+108]\}}{\kappa^4(1+\rho\mu\theta)^4(2+\rho\mu\theta)^4(3+\rho\mu\theta)^3}$
- $\frac{\partial^2}{\partial\theta^2}c_{9,11} = \frac{4\rho\mu^3\theta[3(\rho\mu\theta)^6+9(\rho\mu\theta)^5-43(\rho\mu\theta)^4-201(\rho\mu\theta)^3-212(\rho\mu\theta)^2+36(\rho\mu\theta)+108]}{\kappa^4(1+\rho\mu\theta)^4(2+\rho\mu\theta)^4(3+\rho\mu\theta)^3}$

Table 4: Second-order partial derivatives of $\theta$ to the asymmetric quartet coefficient matrix $\boldsymbol{C}_a$

| $\frac{\partial^2}{\partial\theta^2}\boldsymbol{c}_{a1}$ | $\frac{\partial^2}{\partial\theta^2}\boldsymbol{c}_{a2}$ | $\frac{\partial^2}{\partial\theta^2}\boldsymbol{c}_{a3}$ | $\frac{\partial^2}{\partial\theta^2}\boldsymbol{c}_{a4}$ |
|---|---|---|---|
| $0$ | $0$ | $0$ | $0$ |
| $\frac{2(\kappa-1)\rho^2\mu^2}{\kappa^4(1+\rho\mu\theta)^3}$ | $-\frac{2\rho^2\mu^2}{\kappa^4(1+\rho\mu\theta)^3}$ | $\frac{2(\kappa-1)\rho^2\mu^2}{\kappa^4(1+\rho\mu\theta)^3}$ | $\frac{2(\kappa-1)\rho^2\mu^2}{\kappa^4(1+\rho\mu\theta)^3}$ |
| $\frac{4(\kappa-1)\rho^2\mu^2}{\kappa^4(1+\rho\mu\theta)^3}$ | $\frac{2(\kappa-2)\rho^2\mu^2}{\kappa^4(1+\rho\mu\theta)^3}$ | $-\frac{4\rho^2\mu^2}{\kappa^4(1+\rho\mu\theta)^3}$ | $\frac{4(\kappa-1)\rho^2\mu^2}{\kappa^4(1+\rho\mu\theta)^3}$ |
| $\frac{4(\kappa-2)(\kappa-1)\rho^2\mu^2(3\rho^2\mu^2\theta^2+9\rho\mu\theta+7)}{\kappa^4(1+\rho\mu\theta)^3(2+\rho\mu\theta)^3}$ | $-\frac{4(\kappa-2)\rho^2\mu^2(3\rho^2\mu^2\theta^2+9\rho\mu\theta+7)}{\kappa^4(1+\rho\mu\theta)^3(2+\rho\mu\theta)^3}$ | $-\frac{4(\kappa-2)\rho^2\mu^2(3\rho^2\mu^2\theta^2+9\rho\mu\theta+7)}{\kappa^4(1+\rho\mu\theta)^3(2+\rho\mu\theta)^3}$ | $\frac{4(\kappa-2)(\kappa-1)\rho^2\mu^2(3\rho^2\mu^2\theta^2+9\rho\mu\theta+7)}{\kappa^4(1+\rho\mu\theta)^3(2+\rho\mu\theta)^3}$ |
| $\frac{6(\kappa-1)\rho^2\mu^2}{\kappa^4(1+\rho\mu\theta)^3}$ | $\frac{2(2\kappa-3)\rho^2\mu^2}{\kappa^4(1+\rho\mu\theta)^3}$ | $\frac{2(2\kappa-3)\rho^2\mu^2}{\kappa^4(1+\rho\mu\theta)^3}$ | $-\frac{6\rho^2\mu^2}{\kappa^4(1+\rho\mu\theta)^3}$ |
| $\frac{4(\kappa-2)(\kappa-1)\rho^2\mu^2(3\rho^2\mu^2\theta^2+9\rho\mu\theta+7)}{\kappa^4(1+\rho\mu\theta)^3(2+\rho\mu\theta)^3}$ | $-\frac{4(\kappa-2)\rho^2\mu^2(3\rho^2\mu^2\theta^2+9\rho\mu\theta+7)}{\kappa^4(1+\rho\mu\theta)^3(2+\rho\mu\theta)^3}$ | $\frac{4(\kappa-2)(\kappa-1)\rho^2\mu^2(3\rho^2\mu^2\theta^2+9\rho\mu\theta+7)}{\kappa^4(1+\rho\mu\theta)^3(2+\rho\mu\theta)^3}$ | $-\frac{4(\kappa-2)\rho^2\mu^2(3\rho^2\mu^2\theta^2+9\rho\mu\theta+7)}{\kappa^4(1+\rho\mu\theta)^3(2+\rho\mu\theta)^3}$ |
| $\frac{6(\kappa-1)^2\rho^2\mu^2}{\kappa^4(1+\rho\mu\theta)^4}$ | $-\frac{6(\kappa-1)\rho^2\mu^2}{\kappa^4(1+\rho\mu\theta)^4}$ | $-\frac{6(\kappa-1)\rho^2\mu^2}{\kappa^4(1+\rho\mu\theta)^4}$ | $-\frac{6(\kappa-1)\rho^2\mu^2}{\kappa^4(1+\rho\mu\theta)^4}$ |
| $\frac{8(\kappa-2)(\kappa-1)\rho^2\mu^2(3\rho^2\mu^2\theta^2+9\rho\mu\theta+7)}{\kappa^4(1+\rho\mu\theta)^3(2+\rho\mu\theta)^3}$ | $\frac{4(\kappa-2)^2\rho^2\mu^2(3\rho^2\mu^2\theta^2+9\rho\mu\theta+7)}{\kappa^4(1+\rho\mu\theta)^3(2+\rho\mu\theta)^3}$ | $-\frac{8(\kappa-2)\rho^2\mu^2(3\rho^2\mu^2\theta^2+9\rho\mu\theta+7)}{\kappa^4(1+\rho\mu\theta)^3(2+\rho\mu\theta)^3}$ | $-\frac{8(\kappa-2)\rho^2\mu^2(3\rho^2\mu^2\theta^2+9\rho\mu\theta+7)}{\kappa^4(1+\rho\mu\theta)^3(2+\rho\mu\theta)^3}$ |
| $\frac{8(\kappa-2)^2(\kappa-1)\rho^2\mu^2(6\rho^2\mu^2\theta^2+16\rho\mu\theta+11)}{\kappa^4(1+\rho\mu\theta)^3(2+\rho\mu\theta)^4}$ | $-\frac{8(\kappa-2)^2\rho^2\mu^2(6\rho^2\mu^2\theta^2+16\rho\mu\theta+11)}{\kappa^4(1+\rho\mu\theta)^3(2+\rho\mu\theta)^4}$ | $-\frac{8(\kappa-2)^2\rho^2\mu^2(6\rho^2\mu^2\theta^2+16\rho\mu\theta+11)}{\kappa^4(1+\rho\mu\theta)^3(2+\rho\mu\theta)^4}$ | $-\frac{8(\kappa-2)^2\rho^2\mu^2(6\rho^2\mu^2\theta^2+16\rho\mu\theta+11)}{\kappa^4(1+\rho\mu\theta)^3(2+\rho\mu\theta)^4}$ |
| See $\frac{\partial^2}{\partial\theta^2}c_{9,1}$ | See $\frac{\partial^2}{\partial\theta^2}c_{9,2}$ | See $\frac{\partial^2}{\partial\theta^2}c_{9,3}$ | See $\frac{\partial^2}{\partial\theta^2}c_{9,4}$ |

| $\frac{\partial^2}{\partial\theta^2}\boldsymbol{c}_{a5}$ | $\frac{\partial^2}{\partial\theta^2}\boldsymbol{c}_{a6}$ | $\frac{\partial^2}{\partial\theta^2}\boldsymbol{c}_{a7}$ | $\frac{\partial^2}{\partial\theta^2}\boldsymbol{c}_{a8}$ |
|---|---|---|---|
| $0$ | $0$ | $0$ | $0$ |
| $\frac{2(\kappa-1)\rho^2\mu^2}{\kappa^4(1+\rho\mu\theta)^3}$ | $-\frac{2\rho^2\mu^2}{\kappa^4(1+\rho\mu\theta)^3}$ | $-\frac{2\rho^2\mu^2}{\kappa^4(1+\rho\mu\theta)^3}$ | $\frac{2(\kappa-1)\rho^2\mu^2}{\kappa^4(1+\rho\mu\theta)^3}$ |
| $-\frac{4\rho^2\mu^2}{\kappa^4(1+\rho\mu\theta)^3}$ | $\frac{2(\kappa-2)\rho^2\mu^2}{\kappa^4(1+\rho\mu\theta)^3}$ | $-\frac{4\rho^2\mu^2}{\kappa^4(1+\rho\mu\theta)^3}$ | $-\frac{4\rho^2\mu^2}{\kappa^4(1+\rho\mu\theta)^3}$ |
| $-\frac{4(\kappa-2)\rho^2\mu^2(3\rho^2\mu^2\theta^2+9\rho\mu\theta+7)}{\kappa^4(1+\rho\mu\theta)^3(2+\rho\mu\theta)^3}$ | $-\frac{4(\kappa-2)\rho^2\mu^2(3\rho^2\mu^2\theta^2+9\rho\mu\theta+7)}{\kappa^4(1+\rho\mu\theta)^3(2+\rho\mu\theta)^3}$ | $\frac{8\rho^2\mu^2(3\rho^2\mu^2\theta^2+9\rho\mu\theta+7)}{\kappa^4(1+\rho\mu\theta)^3(2+\rho\mu\theta)^3}$ | $-\frac{4(\kappa-2)\rho^2\mu^2(3\rho^2\mu^2\theta^2+9\rho\mu\theta+7)}{\kappa^4(1+\rho\mu\theta)^3(2+\rho\mu\theta)^3}$ |
| $\frac{2(\kappa-3)\rho^2\mu^2}{\kappa^4(1+\rho\mu\theta)^3}$ | $\frac{2(\kappa-3)\rho^2\mu^2}{\kappa^4(1+\rho\mu\theta)^3}$ | $\frac{2(\kappa-3)\rho^2\mu^2}{\kappa^4(1+\rho\mu\theta)^3}$ | $-\frac{6\rho^2\mu^2}{\kappa^4(1+\rho\mu\theta)^3}$ |
| $-\frac{4(\kappa-2)\rho^2\mu^2(3\rho^2\mu^2\theta^2+9\rho\mu\theta+7)}{\kappa^4(1+\rho\mu\theta)^3(2+\rho\mu\theta)^3}$ | $-\frac{4(\kappa-2)\rho^2\mu^2(3\rho^2\mu^2\theta^2+9\rho\mu\theta+7)}{\kappa^4(1+\rho\mu\theta)^3(2+\rho\mu\theta)^3}$ | $\frac{8\rho^2\mu^2(3\rho^2\mu^2\theta^2+9\rho\mu\theta+7)}{\kappa^4(1+\rho\mu\theta)^3(2+\rho\mu\theta)^3}$ | $-\frac{4(\kappa-2)\rho^2\mu^2(3\rho^2\mu^2\theta^2+9\rho\mu\theta+7)}{\kappa^4(1+\rho\mu\theta)^3(2+\rho\mu\theta)^3}$ |
| $\frac{6(\kappa-1)^2\rho^2\mu^2}{\kappa^4(1+\rho\mu\theta)^4}$ | $\frac{6\rho^2\mu^2}{\kappa^4(1+\rho\mu\theta)^4}$ | $-\frac{6(\kappa-1)\rho^2\mu^2}{\kappa^4(1+\rho\mu\theta)^4}$ | $-\frac{6(\kappa-1)\rho^2\mu^2}{\kappa^4(1+\rho\mu\theta)^4}$ |
| $-\frac{8(\kappa-2)\rho^2\mu^2(3\rho^2\mu^2\theta^2+9\rho\mu\theta+7)}{\kappa^4(1+\rho\mu\theta)^3(2+\rho\mu\theta)^3}$ | $-\frac{8(\kappa-2)\rho^2\mu^2(3\rho^2\mu^2\theta^2+9\rho\mu\theta+7)}{\kappa^4(1+\rho\mu\theta)^3(2+\rho\mu\theta)^3}$ | $-\frac{8(\kappa-2)\rho^2\mu^2(3\rho^2\mu^2\theta^2+9\rho\mu\theta+7)}{\kappa^4(1+\rho\mu\theta)^3(2+\rho\mu\theta)^3}$ | $\frac{16\rho^2\mu^2(3\rho^2\mu^2\theta^2+9\rho\mu\theta+7)}{\kappa^4(1+\rho\mu\theta)^3(2+\rho\mu\theta)^3}$ |
| $-\frac{8(\kappa-2)^2\rho^2\mu^2(6\rho^2\mu^2\theta^2+16\rho\mu\theta+11)}{\kappa^4(1+\rho\mu\theta)^3(2+\rho\mu\theta)^4}$ | $\frac{16(\kappa-2)\rho^2\mu^2(6\rho^2\mu^2\theta^2+16\rho\mu\theta+11)}{\kappa^4(1+\rho\mu\theta)^3(2+\rho\mu\theta)^4}$ | $\frac{16(\kappa-2)\rho^2\mu^2(6\rho^2\mu^2\theta^2+16\rho\mu\theta+11)}{\kappa^4(1+\rho\mu\theta)^3(2+\rho\mu\theta)^4}$ | $\frac{16(\kappa-2)\rho^2\mu^2(6\rho^2\mu^2\theta^2+16\rho\mu\theta+11)}{\kappa^4(1+\rho\mu\theta)^3(2+\rho\mu\theta)^4}$ |
| See $\frac{\partial^2}{\partial\theta^2}c_{9,5}$ | See $\frac{\partial^2}{\partial\theta^2}c_{9,6}$ | See $\frac{\partial^2}{\partial\theta^2}c_{9,7}$ | See $\frac{\partial^2}{\partial\theta^2}c_{9,8}$ |

| $\frac{\partial^2}{\partial\theta^2}\boldsymbol{c}_{a9}$ | $\frac{\partial^2}{\partial\theta^2}\boldsymbol{c}_{a10}$ | $\frac{\partial^2}{\partial\theta^2}\boldsymbol{c}_{a11}$ |
|---|---|---|
| $0$ | $0$ | $0$ |
| $-\frac{2\rho^2\mu^2}{\kappa^4(1+\rho\mu\theta)^3}$ | $-\frac{2\rho^2\mu^2}{\kappa^4(1+\rho\mu\theta)^3}$ | $-\frac{2\rho^2\mu^2}{\kappa^4(1+\rho\mu\theta)^3}$ |
| $-\frac{4\rho^2\mu^2}{\kappa^4(1+\rho\mu\theta)^3}$ | $\frac{2(\kappa-2)\rho^2\mu^2}{\kappa^4(1+\rho\mu\theta)^3}$ | $-\frac{4\rho^2\mu^2}{\kappa^4(1+\rho\mu\theta)^3}$ |
| $\frac{4(\kappa-2)(\kappa-1)\rho^2\mu^2(3\rho^2\mu^2\theta^2+9\rho\mu\theta+7)}{\kappa^4(1+\rho\mu\theta)^3(2+\rho\mu\theta)^3}$ | $-\frac{4(\kappa-2)\rho^2\mu^2(3\rho^2\mu^2\theta^2+9\rho\mu\theta+7)}{\kappa^4(1+\rho\mu\theta)^3(2+\rho\mu\theta)^3}$ | $\frac{4(\kappa-2)(\kappa-1)\rho^2\mu^2(3\rho^2\mu^2\theta^2+9\rho\mu\theta+7)}{\kappa^4(1+\rho\mu\theta)^3(2+\rho\mu\theta)^3}$ |
| $\frac{2(\kappa-3)\rho^2\mu^2}{\kappa^4(1+\rho\mu\theta)^3}$ | $-\frac{6\rho^2\mu^2}{\kappa^4(1+\rho\mu\theta)^3}$ | $-\frac{6\rho^2\mu^2}{\kappa^4(1+\rho\mu\theta)^3}$ |
| $-\frac{4(\kappa-2)\rho^2\mu^2(3\rho^2\mu^2\theta^2+9\rho\mu\theta+7)}{\kappa^4(1+\rho\mu\theta)^3(2+\rho\mu\theta)^3}$ | $\frac{8\rho^2\mu^2(3\rho^2\mu^2\theta^2+9\rho\mu\theta+7)}{\kappa^4(1+\rho\mu\theta)^3(2+\rho\mu\theta)^3}$ | $\frac{8\rho^2\mu^2(3\rho^2\mu^2\theta^2+9\rho\mu\theta+7)}{\kappa^4(1+\rho\mu\theta)^3(2+\rho\mu\theta)^3}$ |
| $\frac{6\rho^2\mu^2}{\kappa^4(1+\rho\mu\theta)^4}$ | $\frac{6\rho^2\mu^2}{\kappa^4(1+\rho\mu\theta)^4}$ | $\frac{6\rho^2\mu^2}{\kappa^4(1+\rho\mu\theta)^4}$ |
| $-\frac{4(\kappa-4)\rho^2\mu^2(3\rho^2\mu^2\theta^2+9\rho\mu\theta+7)}{\kappa^4(1+\rho\mu\theta)^3(2+\rho\mu\theta)^3}$ | $-\frac{4(\kappa-4)\rho^2\mu^2(3\rho^2\mu^2\theta^2+9\rho\mu\theta+7)}{\kappa^4(1+\rho\mu\theta)^3(2+\rho\mu\theta)^3}$ | $\frac{16\rho^2\mu^2(3\rho^2\mu^2\theta^2+9\rho\mu\theta+7)}{\kappa^4(1+\rho\mu\theta)^3(2+\rho\mu\theta)^3}$ |
| $\frac{8(\kappa-4)\rho^2\mu^2(6\rho^2\mu^2\theta^2+16\rho\mu\theta+11)}{\kappa^4(1+\rho\mu\theta)^3(2+\rho\mu\theta)^4}$ | $\frac{8(\kappa-4)\rho^2\mu^2(6\rho^2\mu^2\theta^2+16\rho\mu\theta+11)}{\kappa^4(1+\rho\mu\theta)^3(2+\rho\mu\theta)^4}$ | $-\frac{32\rho^2\mu^2(6\rho^2\mu^2\theta^2+16\rho\mu\theta+11)}{\kappa^4(1+\rho\mu\theta)^3(2+\rho\mu\theta)^4}$ |
| See $\frac{\partial^2}{\partial\theta^2}c_{9,9}$ | See $\frac{\partial^2}{\partial\theta^2}c_{9,10}$ | See $\frac{\partial^2}{\partial\theta^2}c_{9,11}$ |

### B Estimation of the variability and sensitivity matrices

#### B.1 Variability matrix - $J(\boldsymbol{\tau}, \theta)$

The gradient vector of the $N$-taxon species tree composite likelihood is

$$\begin{aligned}
\nabla_{\boldsymbol{\tau},\tilde{\theta}} cl(\boldsymbol{\tau}, \theta | \boldsymbol{n}^D) &= \left( \frac{\partial}{\partial \tau_1} cl(\boldsymbol{\tau}, \theta), ..., \frac{\partial}{\partial \tau_{n-1}} cl(\boldsymbol{\tau}, \theta), \frac{\partial}{\partial \tilde{\theta}} cl(\boldsymbol{\tau}, \theta) \right)^T \\
&= \sum_{j=1}^{\binom{N}{4}} \left( \frac{\partial}{\partial \tau_1} \left[ l(Q_j, \boldsymbol{\tau}^{Q_j}, \theta | \boldsymbol{n}^{Q_j}) \right], ..., \frac{\partial}{\partial \tau_{n-1}} \left[ l(Q_j, \boldsymbol{\tau}^{Q_j}, \theta | \boldsymbol{n}^{Q_j}) \right], \frac{\partial}{\partial \tilde{\theta}} \left[ l(Q_j, \boldsymbol{\tau}^{Q_j}, \theta | \boldsymbol{n}^{Q_j}) \right] \right)^T \\
&= \sum_{j=1}^{\binom{N}{4}} \left[ \nabla_{\boldsymbol{\tau},\tilde{\theta}} \boldsymbol{p}^{Q_j} \right] \cdot \text{diag}(1/\boldsymbol{p}^{Q_j}) \cdot \boldsymbol{n}^{Q_j} \\
&= \sum_{j=1}^{\binom{N}{4}} \boldsymbol{R}_j \boldsymbol{E}_j \boldsymbol{n}^D,
\end{aligned}$$

where $\boldsymbol{R}_j = \left[ \nabla_{\boldsymbol{\tau},\tilde{\theta}} \boldsymbol{p}^{Q_j} \right] \cdot \text{diag}(1/\boldsymbol{p}^{Q_j}) = \left( \frac{\partial}{\partial \tau_1} \boldsymbol{p}^{Q_j} \oslash \boldsymbol{p}^{Q_j}, ..., \frac{\partial}{\partial \tau_{n-1}} \boldsymbol{p}^{Q_j} \oslash \boldsymbol{p}^{Q_j}, \frac{\partial}{\partial \tilde{\theta}} \boldsymbol{p}^{Q_j} \oslash \boldsymbol{p}^{Q_j} \right)^T$ is $N \times 15$ partial derivative matrix and $\boldsymbol{E}_j$ is mapping matrix that maps from site pattern count of dataset $\boldsymbol{n}^D$ to the quartet site pattern count $\boldsymbol{n}^{Q_j}$. Then, the variability matrix can be computed by

$$\begin{aligned}
J(\boldsymbol{\tau}, \theta) &= Var \left[ \nabla_{\boldsymbol{\tau},\tilde{\theta}} cl(\boldsymbol{\tau}, \theta | \boldsymbol{n}^D) \right] \\
&= \left\{ \sum_{j=1}^{\binom{N}{4}} \boldsymbol{R}_j \boldsymbol{E}_j \right\} Var(\boldsymbol{n}^D) \left\{ \sum_{j=1}^{\binom{N}{4}} \boldsymbol{R}_j \boldsymbol{E}_j \right\}^T,
\end{aligned}$$

where the variance matrix can be estimated by the empirical variance $\widehat{Var(\boldsymbol{n}^D)} = [\text{diag}(\boldsymbol{n}^D) - \boldsymbol{n}^D (\boldsymbol{n}^D)^T / L]$ and $L$ is the total length of the DNA matrix.

#### B.2 Sensitivity matrix - $H(\boldsymbol{\tau}, \theta)$

The Hessian matrix of the $N$-taxon species tree composite likelihood is

$$\nabla^2_{\boldsymbol{\tau},\tilde{\theta}} cl(\boldsymbol{\tau}, \theta | \boldsymbol{n}^D) = \begin{pmatrix}
\frac{\partial^2}{\partial \tau_1^2} cl(\boldsymbol{\tau}, \theta) & \cdots & \frac{\partial^2}{\partial \tau_1 \partial \tau_{n-1}} cl(\boldsymbol{\tau}, \theta) & \frac{\partial^2}{\partial \tau_1 \partial \tilde{\theta}} cl(\boldsymbol{\tau}, \theta) \\
\vdots & \ddots & \vdots & \vdots \\
\frac{\partial^2}{\partial \tau_1 \partial \tau_{n-1}} cl(\boldsymbol{\tau}, \theta) & \cdots & \frac{\partial^2}{\partial \tau_{n-1}^2} cl(\boldsymbol{\tau}, \theta) & \frac{\partial^2}{\partial \tau_{n-1} \partial \tilde{\theta}} cl(\boldsymbol{\tau}, \theta) \\
\frac{\partial^2}{\partial \tau_1 \partial \tilde{\theta}} cl(\boldsymbol{\tau}, \theta) & \cdots & \frac{\partial^2}{\partial \tau_{n-1} \partial \tilde{\theta}} cl(\boldsymbol{\tau}, \theta) & \frac{\partial^2}{\partial \tilde{\theta}^2} cl(\boldsymbol{\tau}, \theta)
\end{pmatrix}$$

$$= \sum_{j=1}^{\binom{N}{4}} \begin{pmatrix}
\frac{\partial^2}{\partial \tau_1^2} \left[ l(Q_j, \boldsymbol{\tau}^{Q_j}, \theta | \boldsymbol{n}^{Q_j}) \right] & \cdots & \frac{\partial^2}{\partial \tau_1 \partial \tau_{n-1}} \left[ l(Q_j, \boldsymbol{\tau}^{Q_j}, \theta | \boldsymbol{n}^{Q_j}) \right] & \frac{\partial^2}{\partial \tau_1 \partial \tilde{\theta}} \left[ l(Q_j, \boldsymbol{\tau}^{Q_j}, \theta | \boldsymbol{n}^{Q_j}) \right] \\
\vdots & \ddots & \vdots & \vdots \\
\frac{\partial^2}{\partial \tau_1 \partial \tau_{n-1}} \left[ l(Q_j, \boldsymbol{\tau}^{Q_j}, \theta | \boldsymbol{n}^{Q_j}) \right] & \cdots & \frac{\partial^2}{\partial \tau_{n-1}^2} \left[ l(Q_j, \boldsymbol{\tau}^{Q_j}, \theta | \boldsymbol{n}^{Q_j}) \right] & \frac{\partial^2}{\partial \tau_{n-1} \partial \tilde{\theta}} \left[ l(Q_j, \boldsymbol{\tau}^{Q_j}, \theta | \boldsymbol{n}^{Q_j}) \right] \\
\frac{\partial^2}{\partial \tau_1 \partial \tilde{\theta}} \left[ l(Q_j, \boldsymbol{\tau}^{Q_j}, \theta | \boldsymbol{n}^{Q_j}) \right] & \cdots & \frac{\partial^2}{\partial \tau_{n-1} \partial \tilde{\theta}} \left[ l(Q_j, \boldsymbol{\tau}^{Q_j}, \theta | \boldsymbol{n}^{Q_j}) \right] & \frac{\partial^2}{\partial \tilde{\theta}^2} \left[ l(Q_j, \boldsymbol{\tau}^{Q_j}, \theta | \boldsymbol{n}^{Q_j}) \right]
\end{pmatrix}.$$

The second-order partial derivatives of quartet likelihood with respect to $\boldsymbol{\tau}$ and $\tilde{\theta}$ are shown in A.1.2 and A.2.2. Then the sensitivity matrix can be computed entry by entry.

$$H(\boldsymbol{\tau}, \theta) = -E\left( \nabla^2_{\boldsymbol{\tau},\tilde{\theta}} cl(\boldsymbol{\tau}, \theta | \boldsymbol{n}^D) \right),$$

where the expectation of the quartet site pattern count $E(\boldsymbol{n}^{Q_j})$ in each entry can be estimated by plugging in their empirical count $\boldsymbol{n}^{Q_j}$.

#### C Site pattern sorting algorithm

**Algorithm 1** Quartet site pattern sorting algorithm

**Require:** A quartet site pattern $\boldsymbol{d}^Q$ that includes a combination of four nucleotides
  Count the [*count of the unique nucleotide*] in $\boldsymbol{d}^Q$.
  **if** [*count of the unique nucleotide*]=1 **then** ▷ Must be $xxxx$
    [index of $\boldsymbol{d}^Q$]= 1
  **else if** [*count of the unique nucleotide*]=2 **then**
    Count the [*frequency of the most frequent nucleotide*] in $\boldsymbol{d}^Q$
    **if** [*Frequency of the most frequent nucleotide*]=3 **then** ▷ Could be $xxxy, xxyx, xyxx, yxxx$
      Find the [*location of the least frequent nucleotide*] in $\boldsymbol{d}^Q$
      [index of $\boldsymbol{d}^Q$]= 6-[*location of the least frequent nucleotide*]
    **else if** [*Frequency of the most frequent nucleotide*]=2 **then** ▷ Could be $xyxy, yxxy, xxyy$
      Find the [*location of the second occurrence of the first nucleotide*] in $\boldsymbol{d}^Q$
      **if** [*Location of the second occurrence of the first nucleotide*]=3 **then** ▷ Must be $xyxy$
        [index of $\boldsymbol{d}^Q$]= 6
      **else if** [*Location of the second occurrence of the first nucleotide*]=4 **then** ▷ Must be $yxxy$
        [index of $\boldsymbol{d}^Q$]= 7
      **else** ▷ Must be $xxyy$
        [index of $\boldsymbol{d}^Q$]= 8
      **end if**
    **end if**
  **else if** [*count of the unique nucleotide*]=3 **then**
    Find the [*locations of the most frequent nucleotide*] in $\boldsymbol{d}^Q$
    **if** [*Locations of the most frequent nucleotide*]=(1,3) **then** ▷ Must be $xyxz$
      [index of $\boldsymbol{d}^Q$]= 9
    **else if** [*Locations of the most frequent nucleotide*]=(1,4) **then** ▷ Must be $xyzx$
      [index of $\boldsymbol{d}^Q$]= 10
    **else if** [*Locations of the most frequent nucleotide*]=(2,3) **then** ▷ Must be $yxxz$
      [index of $\boldsymbol{d}^Q$]= 11
    **else if** [*Locations of the most frequent nucleotide*]=(2,4) **then** ▷ Must be $yxzx$
      [index of $\boldsymbol{d}^Q$]= 12
    **else if** [*Locations of the most frequent nucleotide*]=(1,2) **then** ▷ Must be $xxyz$
      [index of $\boldsymbol{d}^Q$]= 13
    **else** ▷ Must be $yzxx$
      [index of $\boldsymbol{d}^Q$]= 14
    **end if**
  **else if** [*count of the unique nucleotide*]=4 **then** ▷ Must be $xyzw$
    [index of $\boldsymbol{d}^Q$]= 15
  **end if**

In this section, we discuss the process of sorting the site pattern counts from DNA data and identifying the category index for a quartet site pattern. To differentiate the site pattern categories of the data set, we identify them by their unique ID code. Note that this is different from the 15 site pattern categories of the quartet subtree, where each category is identified by a numeric index. Inspired by the 15 quartet site pattern categories $\{xxxx, ..., xyzw\}$, we use a similar ID code consisting of letters $\{x, y, z, w\}$ to represent different site pattern categories of the data set because each site pattern consists of only four nucleotides.

The nucleotide appearing most frequently within a site pattern of an $N$-taxon tree is firstly denoted as $x$. The nucleotide with the next highest frequency is then labeled as $y$, followed sequentially by $z$, and the least frequent is lastly labeled as $w$. For example, a site pattern $AAAGGGGG$ is coded as $yyyxxxxx$ because $A$ has less occurrences than $G$. In cases where the nucleotide frequencies are equal in a site pattern, the first nucleotide from the left is labeled as $x$. This ensures that each site pattern is uniquely associated with a single ID code. For another example, $AAGGCCTT$ is coded as $xxyyzzww$. All observed site patterns are identified by their unique ID codes and their counts in the data are aggregated. Note that this vector of counts is denoted by $\boldsymbol{n}^D$, and the corresponding ID codes are kept separately as a matrix, where each row represents a set of ID codes and each column represents a sample of a specific species. We can take a subset of the ID codes matrix by the columns of a quartet subtree, where the rows are the quartet site patterns and they may repeat, and identify the category indices for those quartet site patterns. Then, we can sort the site pattern counts $\boldsymbol{n}^D$ into the 15 category site pattern counts of the $j^{th}$ quartet subtree by $\boldsymbol{n}^{Q_j} = \boldsymbol{E}^{Q_j}\boldsymbol{n}^D$, where $\boldsymbol{E}^{Q_j}$ is a mapping matrix with 15 rows and projects the site pattern counts $\boldsymbol{n}^D$ to the quartet subtree site pattern counts $\boldsymbol{n}^{Q_j}$. Each row of $\boldsymbol{E}^{Q_j}$ is a binary vector that has the same length as $\boldsymbol{n}^D$ and sorts the selected counts of $\boldsymbol{n}^D$ into the corresponding quartet site pattern counts by 15 categories.

To generate the binary vector in each row of $E^{Q_j}$, we need a mechanism that converts the quartet site pattern (in $x$, $y$, $z$, and $w$) to its corresponding category numeric index. First, in a quartet site pattern of four nucleotides, we identify the nucleotides that do not repeat and count the number of unique nucleotides. For example, only the site pattern $xxxx$ has one unique nucleotide, and only the site pattern $xyzw$ has four unique nucleotides. We can identify them by category 1 and category 15, respectively, based on the count of unique nucleotides.

However, the remaining thirteen site patterns will have either two or three unique nucleotides. Within each case, we need another rule to differentiate and identify each site pattern. If the count of unique nucleotides is two, we find the frequency of the most frequent nucleotide, which can be two or three. In the cases of three, the site patterns are $xxxy$, $xxyx$, $xyxx$, and $yxxx$, which correspond to categories 2, 3, 4, and 5, respectively. We observe that the string index of the least frequent nucleotide is identifiable for each site pattern, allowing us to establish a one-to-one mapping between them. For example, $xxyx$ has its least frequent nucleotide $y$ at index 3, which is assigned to category $6 - 3 = 3$. In cases where the most frequent nucleotide is two, the site patterns are $xyxy$, $yxxy$, and $xxyy$, which correspond to categories 6, 7, and 8, respectively. We observe that the string index of the second occurrence of the first nucleotide is identifiable for each site pattern. For example, the first nucleotide for $yxxy$ is $y$ and the index of the second occurrence of $y$ is 4, which is then assigned to category 7.

Finally, if the count of unique nucleotides is three, the site patterns are $xyxz$, $xyzx$, $yxxz$, $yxzx$, $xxyz$, and $yzxx$, which correspond to categories 9, 10, 11, 12, 13, and 14, respectively. We observe that the two-dimensional vector of the string indices of the most frequent nucleotide is identifiable for each site pattern. For example, the most frequent nucleotide for $yxxz$ is $x$ and its indices are $(1, 2)$, which is then assigned to category 11. The pseudocode in Algorithm 1 makes it easy for readers to understand its simplicity, especially since words are cumbersome to represent, and the flow of the sorting mechanism.

### D Priors for tree parameters

Following Rannala et al. (2012), we set prior for the population size parameter as $\theta \sim IG(\alpha_\theta, \beta_\theta)$ when we know a priori $\theta$ is approximately around $E(\theta) = \beta_\theta/(\alpha_\theta - 1)$. Similarly, the prior for root age is $\tau_{n-1} \sim IG(\alpha_{\tau_{n-1}}, \beta_{\tau_{n-1}})$ when we know a priori $\tau_{n-1}$ is approximately around $E(\tau_{n-1}) = \beta_{\tau_{n-1}}/(\alpha_{\tau_{n-1}} - 1)$. In the inverse gamma distribution, we pick the shape parameters $\alpha = 3$ to represent a choice of non-informative prior. Then, we setup prior for other speciation times $\tau_1, ..., \tau_{n-2}$ such that they partition the root age $\tau_{n-1}$ according to a flat Dirichlet distribution.

**Priors for symmetric quartet** For a symmetric quartet, we have $\tau_1$ independent of $\tau_2$ given root age $\tau_3$. Let $x_1 = \tau_1/\tau_3$ and $x_2 = \tau_2/\tau_3$ where $\tau_3$ is given, we have the constraint that $x_1, x_2 \in (0, 1)$ because $\max(\tau_1, \tau_2) \leq \tau_3$. Then conditioning on $\tau_3$, we can put beta prior on $x_1$ and $x_2$ as

$$x_1, x_2 | \tau_3 \overset{iid}{\sim} Beta(1, 1).$$

Then applying the variable transformation $(x_1, x_2) \leftrightarrow (\tau_1, \tau_2)$, we have

$$f(\tau_1, \tau_2 | \tau_3) = \frac{1}{\tau_3^2} \text{ for } 0 \leq \tau_1 \leq \tau_3,\ 0 \leq \tau_2 \leq \tau_3.$$

**Priors for asymmetric quartet** For an asymmetric quartet, we have $\tau_3 > \tau_2 > \tau_1$. Let $x_1 = \frac{\tau_1}{\tau_3}$, $x_2 = \frac{\tau_2 - \tau_1}{\tau_3}$ and $x_3 = \frac{\tau_3 - \tau_2}{\tau_3}$ where $\tau_3$ is given, we can have

$$(x_1, x_2, x_3)|\tau_3 \sim Dir(1, 1, 1).$$

Then applying the variable transformation $(x_1, x_2) \leftrightarrow (\tau_1, \tau_2)$, we have

$$f(\tau_1, \tau_2|\tau_3) = (2!) * \frac{1}{\tau_3^2} \text{ for } 0 \leq \tau_1 \leq \tau_2 \leq \tau_3.$$

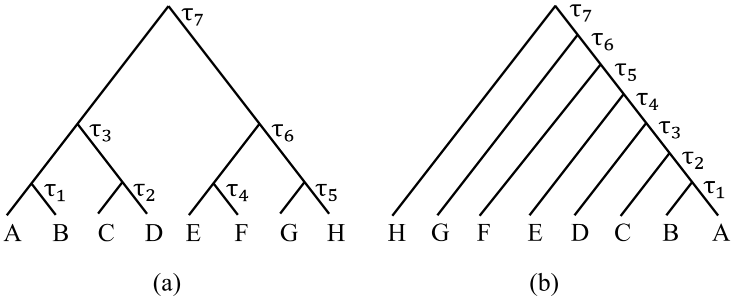


Figure 3: 8 taxa perfectly symmetric and asymmetric tree.

Extending from quartet to full tree with $N$ taxa, we can treat patterns on full tree as either "symmetric" or "asymmetric". In a bifurcating species tree, a "symmetric" pattern arises when both child nodes of an ancestral node are themselves internal nodes. In Figure 3 (a), the root node $\tau_7$ has two child nodes, $\tau_3$ and $\tau_6$, that are internal nodes. So, the nodes of $(\tau_3, \tau_6, \tau_7)$ show a "symmetric" pattern, which is similarly observed in the nodes of $(\tau_1, \tau_2, \tau_3)$ and $(\tau_4, \tau_5, \tau_6)$. Then we have priors $\tau_7 \sim IG(\alpha_\tau, \beta_\tau)$ and

$$\tau_3, \tau_6|\tau_7 \overset{iid}{\sim} Unif(0, \tau_7), \qquad \tau_1, \tau_2|\tau_3 \overset{iid}{\sim} Unif(0, \tau_3), \qquad \tau_4, \tau_5|\tau_6 \overset{iid}{\sim} Unif(0, \tau_6).$$

The prior density function is

$$\pi(\tau_1, ..., \tau_6|\tau_7) = \pi(\tau_1, \tau_2|\tau_3)\pi(\tau_4, \tau_5|\tau_6)\pi(\tau_3, \tau_6|\tau_7) \propto \tau_3^{-2}\tau_6^{-2}\tau_7^{-2}$$

for $\max(\tau_1, \tau_2) \leq \tau_3, \max(\tau_4, \tau_5) \leq \tau_6$, and $\max(\tau_3, \tau_6) \leq \tau_7$.

Another pattern is "asymmetric". For every node in a set of continuous ancestral-descendant nodes, exactly one child node is a leaf and the other an internal node. For example, in Figure 3 (b), the nodes of $(\tau_2, ..., \tau_7)$ are continuous ancestral-descendant nodes and each of them has one child node as a leaf and the other as an internal node. Then, the branch lengths between $\tau_1, \tau_2, ..., \tau_7$ can partition the root age $\tau_7$. We call that the node sequence of $(\tau_1, \tau_2 ..., \tau_7)$ shows an "asymmetric" pattern and we have priors $\tau_7 \sim IG(\alpha_\tau, \beta_\tau)$ and

$$\frac{\tau_2 - \tau_1}{\tau_7}, \frac{\tau_3 - \tau_2}{\tau_7}, ..., \frac{\tau_7 - \tau_6}{\tau_7}|\tau_7 \sim Dir(1, 1, 1, 1, 1, 1).$$

The prior density function after variable transformation is

$$\pi(\tau_1, ..., \tau_6|\tau_7) \propto \tau_7^{-6} \text{ for } \tau_1 \leq ... \leq \tau_6 \leq \tau_7.$$

We conclude this section with a last example in Figure 4. The nodes of $(\tau_5, \tau_6, \tau_7)$ show "symmetric" pattern and we have prior density

$$\pi(\tau_5, \tau_6|\tau_7) \propto \tau_7^{-2}, \text{ for } \max(\tau_5, \tau_6) \leq \tau_7.$$

Similarly for the nodes of $(\tau_1, \tau_2, \tau_3)$, we have prior density

$$\pi(\tau_1, \tau_2|\tau_3) \propto \tau_3^{-2}, \text{ for } \max(\tau_1, \tau_2) \leq \tau_3.$$

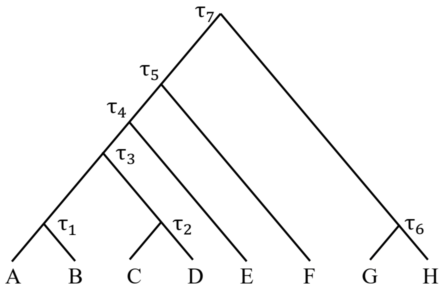

Figure 4: 8 taxa example tree

Since nodes of $(\tau_4, \tau_5)$ are continuous ancestral-descendant nodes with one child node as a leaf and the other as an internal node, we say nodes of $(\tau_3, \tau_4, \tau_5)$ show "asymmetric" pattern. The prior density is

$$\pi(\tau_3, \tau_4 | \tau_5) \propto \tau_5^{-2}, \text{ for } \tau_3 \leq \tau_4 \leq \tau_5.$$

Combining priors above, we have

$$\pi(\tau_1, ..., \tau_6 | \tau_7) \propto \tau_3^{-2} \tau_5^{-2} \tau_7^{-2}, \text{ for } \max(\tau_1, \tau_2) \leq \tau_3 \leq \tau_4 \leq \tau_5, \max(\tau_5, \tau_6) \leq \tau_7.$$
